# RBPSpot: Learning on Appropriate Contextual Information for RBP Binding Sites Discovery

**DOI:** 10.1101/2021.06.07.447370

**Authors:** Nitesh Kumar Sharma, Sagar Gupta, Prakash Kumar, Ashwani Kumar, Upendra Kumar Pradhan, Ravi Shankar

## Abstract

Identifying RBP binding sites and mechanistic factors determining the interactions remain a big challenge. Besides the sparse binding motifs across the RNAs, it also requires a suitable sequence context for binding. The present work describes an approach to detect RBP binding sites while using an ultra-fast BWT/FM-indexing coupled inexact k-mer spectrum search for statistically significant seeds. The seed works as an anchor to evaluate the context and binding potential using flanking region information while leveraging from Deep Feed-forward Neural Network (DNN). Contextual features based on pentamers/dinucloetides which also capture shape and structure properties appeared critical. Contextual CG distribution pattern appeared important. The developed models also got support from MD-simulation studies and the implemented software, RBPSpot, scored consistently high for the considered performance metrics including average accuracy of ∼90% across a large number of validated datasets while maintaining consistency. It clearly outperformed some recently developed tools, including some with much complex deep-learning models, during a highly comprehensive bench-marking process involving three different data-sets and more than 50 RBPs. RBPSpot, has been made freely available, covering most of the human RBPs for which sufficient CLIP-seq data is available (131 RBPs). Besides identifying RBP binding spots across RNAs in human system, it can also be used to build new models by user provided data for any species and any RBP, making it a valuable resource in the area of regulatory system studies.

## Introduction

It has been reported that at any given time, compared to just 2-3% transcription factors expression share, ∼10 times higher volume of RNA binding proteins are expressed (1). Advances with high- throughput techniques like CLIP-seq and Interactome Capture have drastically revised our understanding about RBPs which suggest that human systems are expected to have at least 1,500-2,000 genes coding for RBPs (1, 2). Unfortunately, we are still far behind in terms of information for these regulators where hardly ∼150 RBPs have been studied so far for their interactions with RNAs. Despite of their critical functional roles in cell systems, very few RBPs have been explored with precise identification of their mechanism of action (1).

There are certain limitations with these high-throughput experiments. These experiments are costly. They too don’t give the entire RBP-RNA interactome spectrum and at a time work for one RBP only in condition specific manner. The CLIP-seq reads provide narrowed down regions to look for interactions but don’t provide the mechanistic details and explanations for the interactions. Using general motif discovery tools to identify the interaction spots have got limited success in case of RBPs as they either report too short motifs which have high chances of occurrences across the random data or they don’t cover large spectrum of instances. Unlike transcription factors, RBPs binding sites display sparse motif positional conservation. They are usually difficult to detect through such routine motif finding approaches. Besides the binding motifs, contenxtual sequence environment also guide the RBP-RNA interactions, adding further complexity to the process of discovery of the actual interaction spots. Therefore, this is an area which needs prime focus on deriving the principles of RBP-RNA interactions and their impact of regulation once we have enough CLIP-seq data. One of the most remarkable work, RNAcompete, was done where the authors identified *in-vivo* motifs for 207 different RBPs using pools of 30-41 bases long RNA oligos to which affinity of various RBPs was assessed for binding (3). RNAcompete also highlighted how conventional motif finding tools fail to discover the binding sites motif for RBPs. At computational front some decent progress has been made in dealing with these CLIP-seq data to derive the models for interactions. Initially, to explore the RBPs and their RNA binding sites, databases like RBPDB, CLIPZ, CLIPdb/POSTAR came up (4–7). These databases provided first structured information on RBP-RNA interactions as well as proposed their interaction motifs using traditional motif finding tools while building on publicly available experimental data. As already mentioned above, the motifs being used here are short and occur in abundance even in random data. Also, they don’t consider contextual information. Identification of correct RBP:RNA interaction motifs is a critical step which helps in locating the appropriate contextual information to build an accurate model of RBP:RNA interactions.

RNAcontext is among those first such tools which considered contextual information for RBP-RNA interaction discovery. It applied the structural preferences information for these RNAcompete motifs using *ab-initio* RNA structure prediction tool, sfold (8). However, these *ab-initio* structural prediction methods reliability falls down with the length, making the structural information derived through them not reliable enough (9). The next important stride came with probabilistic tools like RBPmap which extended their previous approach to identify splice sites while applying user provided position specific scoring matrices, supported motif clusters, and phylogenetic conservation to identify RBP RNA interaction spots (10). In the same probabilistic tools category, mCarts was another important addition (11). It works on the similar lines to RBPmap but also applies 6-states Hidden Markov Model (HMM) along with structural information from *ab-initio* secondary structure prediction methods to predicted functional RBP binding sites.

With Graphprot a new generation of such tools started which applied machine learning as well as leveraged from new data-sets developed from CLIP-seq experiments (12). It also applied the concept of differential RNA secondary structure information in contextual manner to build the interaction models. A recently develop tool, beRBP, carries forward the approach similar to RBPmap while implementing a machine-learning method of Random-Forest (13). It clusters the potential motif sites where it ranks them and uses the highest scoring regions for the matches in the given region while scanning for the user provided motif/PWM. In the followup, they have also applied an approach similar to RNAcontext where RNA structural information is provided for the motif region using *ab-initio* structure prediction tool, RNAfold. Further to this, it added the phylogenetic conservation information similar to RBPmap and mCarts.

With recent developments in the area of deep learning, many deep-learning based RBP-RNA interaction detection approaches have been implemented recently. DeepBind deserves special mention among them as it pioneered this category where a robust general system was created to model nucleic acids and protein interactions using convolution neural network (CNN) (14). DeepBind has become a sort of prototype for almost all of the recent Deep-learning based tools to identify the RBP-RNA interactions. DeepBind applies 7-mer motif weight matrices are transformation into an image pixel matrix and is scanned for entire sequence while evaluating for 4- stages to derive the binding score: convolution stage, rectification stage which zooms the scanner to most promising regions for the motif, followed by pooling of all such regions and expansion and clustering of motifs, which is finally subjected to a non-linear classifier. However, the authors accepted that compared to transcription factors and their data, running DeepBind with RNAcompete data did not achieve that level of accuracy. They pointed out the importance of accurate RNA secondary structure information and RNA shape readouts in RNA-RBP interactions which most of the approaches have missed so far. Taking the work further on Deep-learning based RBP-RNA interaction detection, another prominent tool system is iDEEP which has come like a series of softwares like iDeep, iDeepS, and iDeepE (15–17). These tools differ from each other for the way they applied various combinations of CNN and RNN layers. iDeepS applied CNN with Long-Short term memory (LSTM) while taking input from sequence and RNAshape data. iDeepE applies combinations of CNNs which capture local and global sequence properties. A recently developed tool, DeepRiPe, has evolved a CNN and GRU based deep-learning approach while also introducing transcript’s regions specific information like splice junctions etc (18). DeepCLIP is another recently developed tool which detects RBP-RNA interaction spots while applying CNN in combination with bidirectional-LSTM and claims to detect sequence position specific importance which could determine the contribution of various nucleotides in RBP binding (19). These very recently developed deep-learning approaches have become much more complex than DeepBind and claim to achieve much higher accuracy. Their complexity comes from adding complex layers above the regular dense hidden layer. These complex layers actually do the job of automatic feature extraction unlike the other machine-learning approaches where expert knowledge is applied to identify the important properties to look into for feature extraction.

While reviewing these developments and tools, it looked imminent that there is an enormous scope of improvement in the approaches to find and locate RBP-RNA interaction spots. Some of the major points to consider would be: 1) Choice of datasets: A notable issue with all these algorithms is the choice of data-sets, especially the negative data-sets, which have mostly been too relaxed and unrealistic, due to which these tools are prone to over-fitting and imbalance. They are either randomly shuffled sequences or regions randomly selected from those RNAs which did not bind the given RBP. 2) Motif searching approach: most of existing tools, with exception of recent deep- learning based approaches, begin with predefined/user defined motif or PWM derived from traditional motif finding tools with user defined length, which is not a natural approach and one of the prime mistakes. RBP binding sites display sparse conservation which regular motif discovery tools may fail to capture sufficiently. Third, high dependence on *ab-initio* RNA structure prediction tools to derive the structural and accessibility information may be misleading, as already pointed out above, such tools don’t provide correct information on actual complete RNA length. A better approach has been consideration of dinucleotide densities for such purpose (20, 21). Consideration of RNA-shape appears very much important as pointed out by DeepBind as well as some other recent works (14,22,23). It has been reported that pentamers capture the essence of nucleic acid’s shape accurately (24), making them a suitable candidate to be evaluated along with dinucleotide densities to derive RNA structure and shape information. Fourth, though the recent deep-learning approach claim good success through automation of the process of feature extraction at the cost of added complexity, the effectiveness of such automated feature detection needs to be evaluated. Simpler models, if trained with carefully selected properties, are capable to outperform complex models. This is why some of the shallow learning methods have outperformed deep-learning methods on structured data (25, https://towardsdatascience.com/the-unreasonable-ineffectiveness-of-deep-learning-on-tabular-data-fd784ea29c33).

Considering these all factors, here we present a reliable Deep Neural Net (DNN) based approach to build the mechanistic models of RBP-RNA interactions using high-throughput cross-linking data while considering data from 99 experiments and for 137 RBPs for human system. An ultrafast k- mer spectrum search approach was used to identify the most important seed regions in the sequence for contextual information derivation. Contextual information for 75 bases flanking regions around the identified seed derived motif was extracted in the form of variable windowed position specific dinucleotide, pentamers, and heptamers density based propensities. The combined contextual information was provided to a two hidden layers based dense feed-forward networks to accurately identify the RBP binding spots in RNAs. The developed models were used to identify the interaction spots and scored very high accuracy with remarkable balance between sensitivity and specificity as well as performance consistency when tested across a large number and different types of experimental datasets. Molecular dynamics studies also supported these models. The developed approach has been implemented as a freely available webserver and standalone software, RBPSpot. It was comprehensively bench-marked across three totally unbiased standardized data- sets for performance along with five recently published tools, including more complex deep- learning based tools, where it outperformed all of them consistently across all these datasets for most of the studied RBPs. Unlike most of the existing software which don’t provide the option to build new models from data, RBPSpot approach can be applied to detect human system RBP-RNA interactions with its inbuilt models as well as it can be used to develop new models for other species and new RBP data also.

## Materials & Methods

### Data retrieval and processing

The study has considered human RBP models while using high-throughput sequencing data from cross-linking experiments using various CLIP-seq techniques like CLASH, dCLIP, eCLIP, FLASH- CLIP-seq, HITS-CLIP, iCLIP, PAR-CLIP, sCLIP-seq, uvCLAP-CLIP-seq. This data also includes the two cell lines eCLIP data from ENCODE. Most of them are processed peak data collected for 137 RBPs with starBase 2.0 as their primary source (26). A total of 872Mb peak data from 99 experiments were covered in this study for RBP-RNA interaction information from CLIP-seq experiments (Supplementary Data 1 Sheet 1). The peak data of RBPs were downloaded in the form of co-ordinates along with their associated RNA information on which they were binding. Peak data were converted into BED file format along with their strand specificity information. Genome sequences of human hg19 builds were obtained from UCSC browser. Peak data were also refined based on the length distribution and peaks laying in extreme range (length >300 bases and <5 bases) were omitted from the study (Supplementary Data 1 Sheet 2).

### Identification of motif seed candidates: k-mer spectrum search using BWT/FM-Indexing

To search binding sites motifs/seeds for any particular RBP, all the peak regions were transformed into overlapping lists of k-mers of size six to start with. Iteratively and in parallel these generated k- mer spectrum for each such sequence was searched across all the reported cross-linked associated regions in the targets to obtain the enrichment status of the k-mers (seeds) on which motif would be built. These searches were allowed with maximum 30% mismatches. Since normal search would be heavily time consuming step, we implemented an enhanced Burrow-Wheeler transformation with FM-Indexing to search with any number of mismatch which made the search ultra-fast for even in- exact searches. The detailed algorithmic implementation pseudo-code of the implemented algorithm is given in the supplementary methods.

### Identification of motif seeds candidates: Anchoring with the significant seeds

All the k-mer seeds and their relatives displaying at least 70% similarity were evaluated for existence across at least 70% of peak data. Such motif seed candidates were further evaluated for their statistical significance. Those RBPs where no k-mer and their relatives crossed 70% representation were looked for the highest representation available. The remaining data which did not show the representative k-mer were checked further and recursively with minimum cut-off of 20% data representation. Motifs coming from such data were considered as mutually exclusive one. Null model distribution probabilities of occurrence of each k-mer along with its relatives was calculated from the random data set to find their random probabilities. Random data set was generated from unassociated RNAs while randomly carving out the lengths similar to the peak data.

Significantly over-represented k-mers were screened using binomial test with p-value cut-off of 0.01. These significantly enriched k-mers were used as initial seeds to develop the final motif. These seeds of significantly enriched k-mers were expanded in both the directions by expanding by one nucleotide both sides, followed by search across the peak data with at least 70% occurrence in the peak data while repeating the above mentioned search operation recursively. Expansion of seed region in both directions was allowed till at least 70% match existed. Final motifs were selected on the basis of satisfying both the criteria i.e. the motif displayed least 70% abundance across the CLIP-seq instances at 1% significance level and the maximum k-mer expansion maintained at least 70% identity with the associated sequences and relatives. Mutually exclusive motifs were other predominant motifs which existed in the remaining data which were scanned in similar recursive manner as described above. Figure 1. shows the part of the k-mer based motif seed discovery and steps taken afterwards. (Supplementary Data 1 Sheet 3,4)

**Figure 1:**
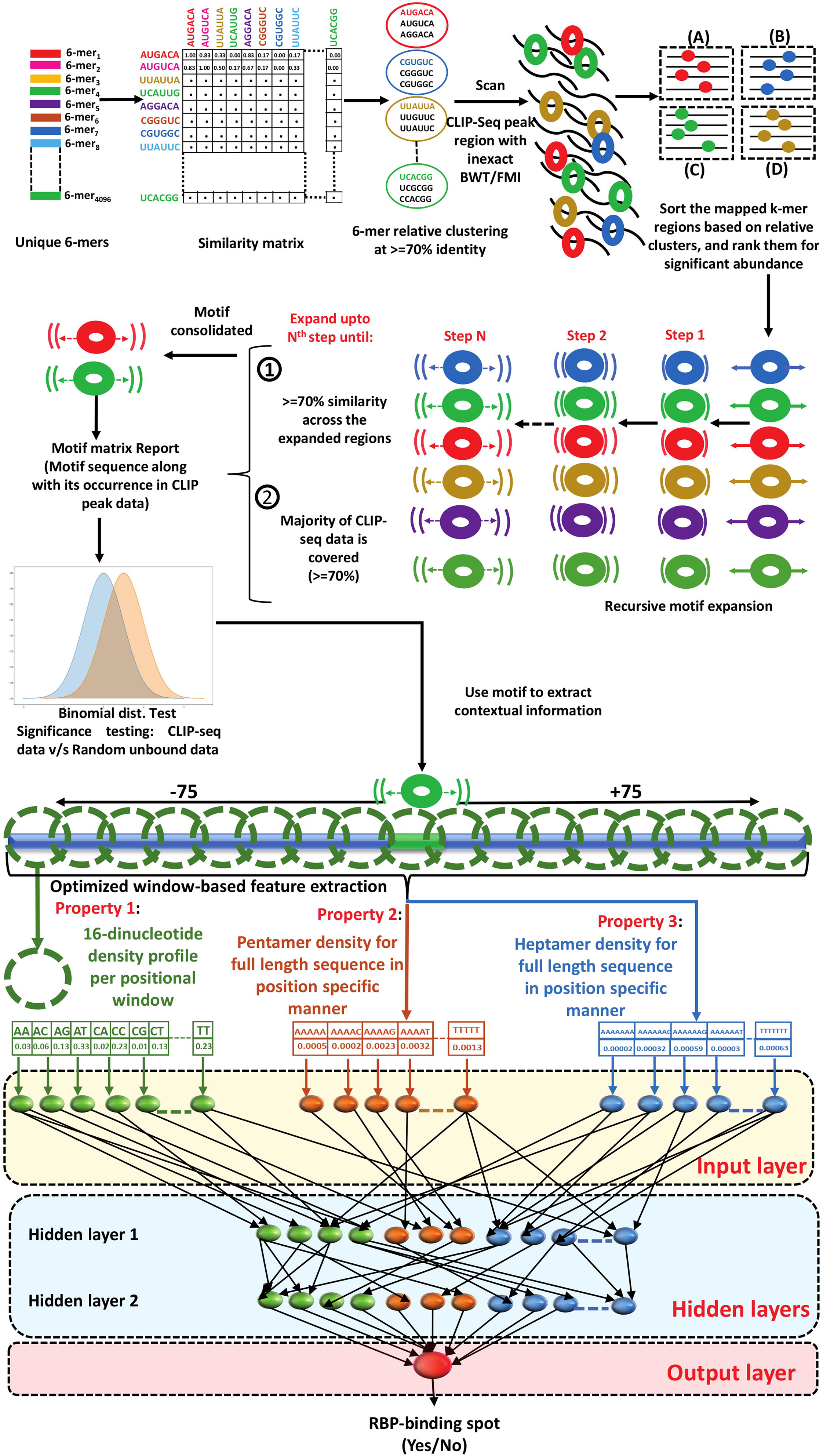
Detailed pipeline of the workflow. The image provides the brief outline of entire computation protocol implemented to develop accurate RBP RNA interaction model and identify the correct RBP binding sites across given RNA sequences. The process of model building starts from identifying prime motifs through k-mer spectrum search from the CLIP-seq regions where BWT/FM indexing based inexact search algorithm was implemented. The statistically enriched k- mers were expanded across all reporting sequences region till at least 70% similarity between them was present. The final prime motifs were established as the anchors. The flanking regions around such anchored prime motifs were used to derive the contextual information, together which worked as freature vector elements for discrimination using XGBoost and Deep-Learning.

### Datasets creation

Once we had prime motifs anchored for each RBP from the given data, their associated peak data sequences were converted into positive datasets. To generate positive datasets for each RBP, start and end co-ordinates from the main motif’s both terminals were expanded by +75 and -75 bases into both the directions. In case of multiple motif locations originating from a single peak for the main motif, all the locations were expanded. Different length dataset sequences formed for different RBPs which depended mainly upon the length of the core motif region. However, for any single RBP all the sequences of the dataset were of same length.

To generate the negative datasets for each RBP, similar condition corresponding RNA-seq data were downloaded from GEO. With minimum three replicates of RNA-seq data expression of each RNA was calculated. Only those transcript sequences were considered which had expression condition available for the same condition but did not bind the RNA or which was not found present in the corresponding condition’s CLIP-seq binding data. Associated main motifs for the RBP were searched across these RNA sets also just in the similar manner as was done to the positive dataset instances. Locations of the main motif were reported in the form of start and end co-ordinates from where further expansion of +75 and -75 bases was done on both the sides. This way very strong negative data-sets were built which ensured that learning was in no way influenced by the motif alone as the motif may also occur randomly to some extent and surrounding context is also considered along in a right manner. This approach was carried out for 74 RBPs for which similar condition RNA-seq data were available. Datasets derived this way were called Set A data-sets.

For 57 RBP similar RNA-seq data were not available for the corresponding conditions. In such scenario the main motifs for negative datasets were searched in those regions which did not appear in the CLIP-seq data but belonged to the same target RNA sequences whose some part appeared in the CLIP-seq, suggesting that though the RNA expressed and even bound to the RBP, these regions despite of having the motif for the RBP did not bind to the RBP and may work as a suitable negative dataset. +75 and -75 flanking bases from both the terminals of the motifs were considered along with the motif region to build the negative datasets. These data-sets were called Set B data- sets.

### Feature generation for positive and negative datasets

Five different types of properties were considered for input into machine learning: 1) The main motif itself, 2) Di-nucleotide density in the associated region while considering 75 bases flanking regions from both the sides of the motif, 3) Dot bracket representation of the RNA structural triplet for the data-set sequences, covering twenty seven combinations of structure triplets arising from the dot-bracket structural representation from RNAfold predicted RNA structures [.((,.(),.(.,.) (,.)),.).,..(,..), …, (((, ((), ((., ()(, ()), ()., (.(, (.), (..,)((,)(),)(.,))(,))),)).,).(,).),)..,], 4) Pentamers density profile for each position which captures the shape information, and, 5) Heptamers densities for the complete region. Dinucleotide densities were evaluated for their discriminatory power for multiple sliding windows starting from 17 to 131. Similarly, the dot brackets structural triplets representation of the data-set sequences were generated using RNAfold (27). They too were evaluated for optimum windows size while testing for window sizes ranging from 29 to full sequence. 1,024 pentamers and 16,384 heptamers densities were evaluated in the similar manner across the data-set sequences.

To calculate heptamers based feature, all positive datasets were split into k-mers of seven bases. Probability of each k-mer were calculated with maximum of two mismatches for each position and accordingly populated in the tensor. Thus, we had 16,384 X ((sequence length) - 7) tensor of probabilities. 16,384 rows represent the heptamers and 150 columns represent individual positions. In the similar manner pentamer features were calculated. For that we had 1024 X ((sequence length)-5) tensor of probabilities. These both tensors were used to convert the sequence data into vectors of probabilities. All together, based on optimum windows, the combined features sets representation of all the data-set sequences was done. The optimum windows and total features varied for each RBP. Finally, each data-set was broken into training and testing data-sets ensuring that no instances from training ever appeared in the testing data-sets. The breakup for each RBP for their training and testing data-sets is given in Supplementary Data 1 Sheet 5 and 6.

### Features evaluation on data-sets

After generating all the features from positive and negative data-sets these features were individually checked for their performance using tree based approaches which are expected to perform better on high dimension instances. Random forest and XGBoost were applied. Each property and their associated feature sets were evaluated for the varying window sizes for their discrimination power between the positive and negative sets. Sliding windows of variable sizes were used for dinculeotide and structure based features. These variable sizes windows were evaluated for the performance. Out of these different sized windows the size producing the best performance was kept for final model generation. It was found that the best performing window size varied across the RBPs, resulting into different optimum windows for the RBPs.

Pentamers and heptamers appeared most informative on the full length window. Equal number of positive and negative instances were chosen for all RBPs considered in the study. From the total chosen instances, 60% were used to create the training set, while remaining 40% instances were used to create the testing set. Python scikit-learn library was used for the same purpose. For feature importance evaluation F-score was used for every considered feature. F-score locates the features which display major difference between their values between negative and positive training sets while comparing the averages for the feature values for positive, negative, and whole set of instances (28). The F-score is represented by the following equation:

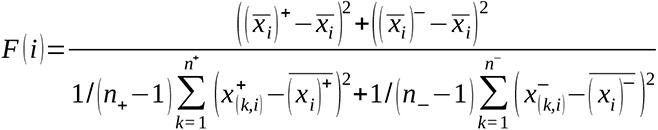

Where:

F(*i*) = Feature score for the ith feature,

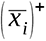= Avergae for i-th feature across the positive instances

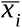 = Total average of the i-th feature across the complete data-set

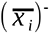= Avergae for i-th feature across the negative instances

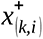 = Feature value for *k-th* instance for *i-th* feature in positive data-set

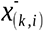 = Feature value for *k-th* instance for *i-th* feature in negative data-set

*n_+_* _=_ Total number of positive instances

*n_-_* = Total number of negative instances

Also, for every *i-th* feature, t-test was conducted between *n_+_ and n_-_* to evaluate the significance of *i- th* feature for its discrimination capability between positive and negative instances.

### Machine learning implementation

With the optimized windows in the above mentioned section, feature vectors for all the RBPs were used to build models to recognize RBP binding sites using two major machine-learning approaches: XGBoost and Two Hidden Layers based Deep Feed Forward Neural Networks (DNNs). Both were implemented using python scikit-learn, Keras, and Tensorflow libraries. In both the cases 70% and 30% of data were retained for train and test sets, respectively.

The DNNs were built where the input layers had number of nodes equal to the number of features for the RBP considered. Thus, the size of input layer varied from 1,200 to 2,500. The performance of DNN was also evaluated for various numbers of hidden layers where finally total two hidden layers were found performing the best. The connections between the nodes were made dense. For every RBP model the number of nodes across the two hidden layers varied between 700 to 1,300. Different types of activation functions combinations were applied for the layers from a pool of a number of available activation functions. Activation functions define the layers and transform the activation values obtained from previous layer to a non-linear form, creating several hyperplanes to obtain best possible discrimination of instances. In most of the models here, the first hidden layer had RELU and the second hidden layer had ELU (for some cases they interchanged also), while the final output layer had sigmoid function.

Every learning step provides estimation of error made, measuring the error and accordingly corrections in the learning rate and weights on connections are done. This error estimation is achieved by loss/cost functions. Multiple types of loss functions were tried to optimize the accuracy. The best performance was obtained for Binary Cross Entropy. Since its a feed forward network where the cost function assess the missed targets and accordingly network connection weights are updated though some optimizer. The optimizer parameter which worked the best was ‘Adam’ optimizer, otherwise SGD with momentum. Usually Adam optimizer works better because of its capability to provide different learning rates per parameter, deals better with sparse gradients, and adapts based on recent learning rates while keeping them in memory. Momentum was applied in the learning which helps to ward-off entrapment under local minima during the minimization steps. The learning rate varied from 0.001 to 0.01 and momentum varied from 0.05 to 0.9. L1 and L2 weight decay regularizors were applied to avoid over-fitting. DNN models were trained using 1000 epochs and batch sizes varying from 50 to 200 instances. All the model from DNN and XGBoost were saved in protobuf format. Since the entire system is implemented here using TensorFlow, the protbuf file provides the graph definition and weights of the model to the TensorFlow structure. The optimum parameter values were fixed using an in-house developed script which tested various combinations of values of the paramters to pick the best ones.

In XGBoost, grid search was applied for parameter optimization. Following parameters were finalized after the grid search: params = {“eta/learning rate”: 0.2, “max_depth”: 4, “objective”: “binary:logistic”, “silent”: 1, “base_score”: np.mean(yt), ’n_estimators’: 1000, “eval_metric”: “logloss”}. Gradient boosted decision trees learn very quickly and may overfit. To overcome this shrinkage was used which slows down the the learning rate of gradient boosting models. Size of the decision tree were run on max-depth=9. At the value of 4 stability was gained as the logloss got stabilized and did not change thereafter.

To evaluate the consistency of performance models developed with the given features, 10-fold cross validation was also performed for each RBP. Everytime, the training dataset was split into 70:30 ratio with first used to train and second part used to test, respectively. Each time data was shuffled and random data was selected for building new model from scratch. This process was repeated 10 times for each RBP. Accuracy and other perfomance measure were calculated for each model. (Supplementary data 1 sheet 7)

The performance on test sets was also evaluated. Confusion matrices containing correctly and incorrectly identified test set instances were built for each RBPs. Frequently used measures for classifier performance evaluation and accuracy of RBPs models were evaluated. Sensitivity (Sn)/Recall/True Positive Rate (TPR) defines the portion of positives which were correctly identified as positives whereas specificity describes the portion of negative instances correctly identified. Precision estimates the proportion of positives with respect to total true and false positives. F1-score was also evaluated which measures the balance between precision and recall. AUC/ROC were also measured for each model. Besides these metrics, Mathew’s Correlation

Coefficient (MCC) was also considered. MCC is considered among the best metrics to fathom the performance where score equally influenced by all the four confusion matrix classes (true positives, false negatives, true negatives, and false positives) (29). A good MCC score is an indicator of robust and balanced model with high degree of performance consistency.

Performance measures were done using the following equations:

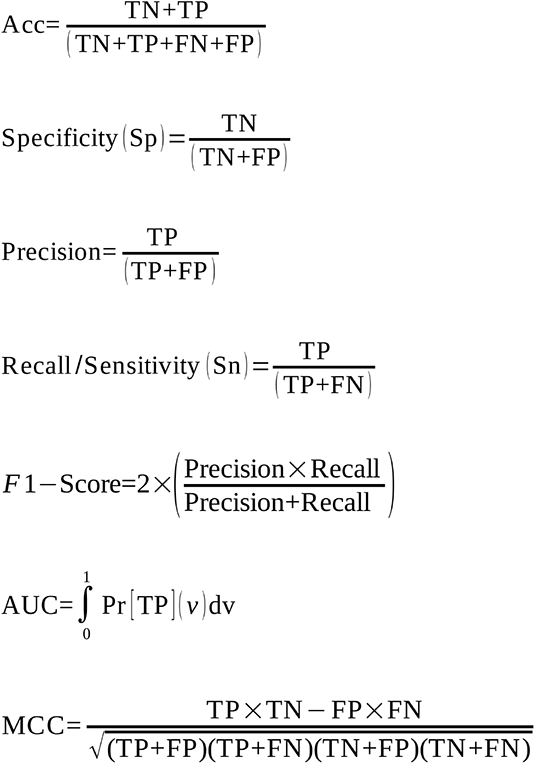

Where:

TP = True Positives, TN = True Negatives, FP = False Positives, FN = False Negatives, Acc = Accuracy, AUC = Area Under Curve

### Structural analysis of identified binding spots

To assess the stability and dynamics of the RBP-RNA complexes for the identified binding spots, structural analysis was done. The 3D coordinates of RBPs were retrieved from the Protein Data Bank (PDB). X-Ray crystallographic structure for 13 different RBPs were downloaded. Prior to docking, protein structures were prepared by removing water molecules and other hetero-atoms, while adding polar hydrogen atoms. RNA motifs identified through RBPSpot algorithm for above mentioned five RBPs were taken as flexible molecules. All docking studies were performed through NPDock (Nucleic Acid–Protein Docking) and PATCHDOCK incorporating more realistic DARS-RNP statistical potential based on reverse Boltzmann statistics to score protein-RNA complexes (30). RNA motifs three dimensional structures were built using RNACOMPOSER web server based on RNA FRABASE database relating the RNA secondary and tertiary structure elements. In order to search for all possible RNA-binding sites and optimize the structural effects of RNA on the construction of complex, short RNA motifs were taken into account. Protein-RNA interface residues were predicted using DR_Bind1 (31) based on evolutionary conservation. Top three representative docking potential-ranked protein-RNA complexes were built for each of the above mentioned RBPs and the best one was considered for further analysis.

### MD simulations

All molecular dynamics simulations of the RBP alone and the RBP–RNA complex were conducted using GROMACS 5.1 package (32), modeling each system with the AMBER03 force-field of protein and nucleic acids (33) with periodic boundary conditions. The topology files for the selected target RNA motifs were built using pdb2gmx in the framework of AMBER03 force-field. Models were solvated with the TIP3P water model (34). The distance between the biomolecule and the edge of the simulation box was set as minimum 1.0 Å so that they could not directly interact with their own periodic boundary condition and fully immerse with water while rotating freely. Boxes were solvated with TIP3P water. The number of solvated molecules added to each system varied. After the establishment of initial configuration, the systems were minimized. 50,000 steps (steepest descent approach) were used for each system until the maximum force of < 10.0 kJ/mol for energy minimization. For calculation of long range electrostatic interactions, Particle Mesh Ewald (PME) method was used. To establish the systems at constant temperature of 300K, V-rescale thermostat (modified Berendsen thermostat), at a constant pressure of 1 bar, and Parrinello-Rahman berostat were applied with a 2 ps coupling constant for both parameters. The LINCS algorithm (35) was used to constrain all bond lengths involving hydrogens. During the production run, a time step of 2 fs was used and conformations were saved every 10 ps for the analysis of molecular dynamics trajectory of total 20 ns for each RBP and their complexes using leap-frog algorithm (36) to integrate the equation of motion. MD trajectories were further evaluated for considering Root Mean Square Deviation (RMSD). RMSD is suitable to decipher the structural changes in proteins and their complex structures corresponding to initial structure during the course of different time periods of dynamics simulation. RMSD was calculated using the following equation:

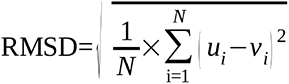

where,

u_i_=Cartesian coordinates of atom *i* in the initial structure;

v_i_=Cartesian coordinates of atom *i* in the structure during simulation;

N=number of atoms;

To analyze the structural properties of the individual RBPs and their complexes in the form of root mean square deviation (RMSD), g_rms functions were utilized. Changes in trajectories of molecular dynamics during course of simulation were plotted for evaluation using python plotting library.

### Co-occurring RNA motifs group clustering

A two steps statistical approach was employed to identify the co-occurring motif pairs. In this approach, the positive set of RBP was scanned for other most frequent occurring k-mers. Top co- occurring motifs were checked for their statistical significance. KS-test was used to find the significance of distance for two motifs. All the distance between two motifs were calculated from positive and negative data-sets. Distribution plot of random data and positive data were further checked using KS-test. Level of significance were considered p<0.05. They were further checked for frequency ratio (FR). At 5% level of significance, if the hypergeometric test *p-value* was less than 0.05, motif pair of enriched and co-occurring motifs was considered significant. Additionally, frequency ratio (FR) as a measure of co-occurrence of motif pairs was also computed to estimate the tendency of motif pairs to co-occur with each other as proposed previously (37):

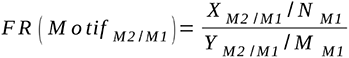

X_M2/M1_=Number of sequences containing motif1

N_M1_=Number of sequences containing motif2 co-occurring with motif1

Y_M2/M1_=Number of sequences without motif1

M_M1_=Number of sequences containing motif2 without motif

### Benchmarking and Performance Evaluation

To evaluate the RBPSpot performance and the importance of dataset constructed in this study, we compared RBPSpot with five different tool: RBPmap, DeepBind, iDeepE, DeepCLIP, and beRBP. Three different datasets were considered separately for the benchmarking process: Datasets used for RBPSpot, beRBP, and Graphprot. Datasets of beRBP and Graphprot are common data source for most of the existing published software built to identify RBP-RNA interactions. As already mentioned above, RBPSpot dataset is based on the positive datasets from ENCORI (the encyclopedia of RNA Interactomes, previously known as StarBase) and the negative datasets based on the protocol mentioned above in the previous section. This dataset contained positive and negative sequences for 131 RBPs in which length of sequence varied from minimum of 156 to maximum of 160 bases. The variation in the length of the sequences for different RBPs was due to the varying length of their major motifs. For benchmarking purpose those RBPs data were considered from this dataset for which at least one tool had model ready for comparison. No such RBP was considered from this dataset for benchmarking for which no other tool had model ready for comparison. This way a total of 52 RBP data were used from RBPSpot dataset for the comparison purpose.

The beRBP dataset is available for 29 RBPs. This dataset is based on the experimentally validated target sequences (3′-UTRs) for human RBPs (positive datasets) from AURA (38) (v2, 8/5/2015;http://aura.science.unitn.it/), which is a manually curated and comprehensive catalog of human UTRs bound by regulators including RBPs. Negative instances of this dataset has random sequences chosen from the 3′-UTR pool. The beRBP dataset was obtained from the URL http://bioinfo.vanderbilt.edu/beRBP/download/TabS1.7z.

The third dataset considered in this study was built during the work presenting Graphprot software. Since then, this dataset has been used extensively by many published software to this date. This dataset covers 24 RBPs coming from various CLIP-seq experiments. For each set of CLIP-seq data, they created a set of unbound sites by shuffling the co-ordinates of bound sites within all genes occupied by at least one binding site which worked as the negative dataset. making the corresponding negative dataset instances. This dataset was retrieved from URL http://www.bioinf.uni-freiburg.de/Software/GraphProt/GraphProt_CLIP_sequences.tar.bz2.

The four out of the compared five tools *viz.* beRBP, RBPmap, DeepCLIP, and DeepBind provide pre-built models. Only iDeepE does not provide any pre-built model. To overcome this, models were generated using iDeepE methodology for the datasets. To make binary decisions with DeepBind, threshold of 0.7 was applied after performing logistic transformation of the raw DeepBind scores (39).

In the second part of the benchmarking impact of datasets was assessed on model building quality where models were built using different datasets and various comabinations of test and train datasets were analysed. Besides RBPSpot, only two tool, iDeepE and DeepCLIP, had provision to build models from user provided datasets. Remaining tools have fixed models with which they work and don’t provide the provision to build models from user provided data. Therefore, they could not be included in this part of benchmarking. Thus, ror this part, the datasets used by RBPSpot (RBPSpot dataset), iDeepE, and DeepCLIP (Graphprot dataset) were used. Four differenet combinations of train and test datasets (RBPSpot train and RBPSpot test, RBPSpot train and Graphprot test, Graphprot train and RBPSpot test and Graphprot train and Graphprot test) were used for the benchmarking to evaluate the impact of datasets on the performance of these algorithms.

### Comparison with experimentally reported motifs

A total of 29 RBPs from RNAcompete study were found overlapping with our set of 131 RBPs. Their IUPAC motifs were downloaded from RNAcompete web portal. For these 29 RBPs a total of 44 motifs were reported. Out of these 44 motifs, 35 motifs had a length of 7 bases, eight motifs had a length of 6 bases, and one motif had a length of 5 bases. Four motifs out of 44, were discarded due to more than 3 variable positions in a length of 7 bases. Therefore, in the final analysis a total of 40 motifs representing 26 RBPs, were present. These motifs were scanned in the similar manner as was done with the search for motifs identified by RBPSpot approach in order to maintain an unbiased motif search approach. Random data sets to evaluate the random chance observations were generated from the transcriptome data using the length exactly similar to the ones from the cross- linking peak data. The similar above mentioned allowed mismatches based motif searching criteria was used here also to scan the random datasets for motif occurrence in them. Binomial test was applied to find the significance of these motifs in the cross-linking data. Other than RNAcompete motif, experimentally validated motifs were also considered from CISBP-RNA Database. A total of 31 RBPs from this dataset were found overlapping with RBPSpot data. Out of these 31 RBPs, 24 RBPs were reported from RNACompete study only, two RBPs were reported through SELEX and yeast three-hybrid screening whereas five RBPs were reported from RNAcompete and SELEX/RIP- Chip. These motifs were also searched in the similar manner.

### Application of RBPSpot across SARS-CoV2 genome

To identify the binding sites of RBPs across SARS-Cov2 genome, we downloaded its genome from NCBI (accession number NC_045512).

## Results and Discussion

### Reads data collection, filtering, and pre-processing

CLIP-seq peak data from various sources were collected for 137 RBPs from starBase 2.0, also know as ENCORI (Supplementary Data 1 Sheet 1). All the data were collected in the form of co- ordinates. These data were from multiple types of CLIP-seq experimental techniques i.e. CLASH, dCLIP, eCLIP, FLASH-CLIP-seq, HITS-CLIP, iCLIP, PAR-CLIP, sCLIP-seq, and uvCLAP. The peak data varied from 234 (PAPD5) to 9,84,503 (U2AF2) peaks. Initially, six RBPs’ data were discarded due to insufficient peak data availability. Here we considered only those RBPs which were having >500 unique binding peaks available. These six RBPs *viz.* PAPD5 (234), EIF3B (298), EIF3A (371), EIF3G (398), EIF3D (399), and PUM1 (473) had lesser number of initial peaks available. Remaining data for 131 RBPs were having a total number of 2,11,23,594 unique peaks. To further filter this data we discarded those sequences which were having a length <5 nucleotides or extreme length sequences (>300 basepairs). With this all, a total of 1,87,14,999 peaks were available for the study, varying from EIF4A1 (1, 175) to AGO1-4 (9,41,224). Initial co-ordinate data were extracted into sequences from genome. Initial and final data are given in (Supplementary Data 1 Sheet 2).

### Most of the RBP binding sites display a prime binding motif covering majority and along with co-occuring motifs

As discussed in the introduction section, most of the available tools for identifying the RBP bindings sites across the RNAs require either prior information available traditional motif finding approaches like MEME, TOMTOM or HOMER. The application of traditional motif discovery tools may not be much information in case of RBPs which have been reported to be sparse, short, and poorly conserved. Further to this, such motif discovery approaches expect user defined motif length instead of naturally capturing the motif. In general, if such motifs are not considered with proper context they may lead towards false discoveries. Here, we have used the initial deep-sequencing data to find the most frequently occurring k-mers (seeds) to make it an initial step for motif finding. To find a naturally occurring most frequent k-mers, search was started with k=6 with two mismatches. Reason behind this was that some of the previously reported motifs for RBPs were either very sparse or as small as 4 bases long only. Therefore, a k-mer with six bases and with two mismatches would fetch all possible 6-mer spectrum which would agree with each other with two mismatches (relatives to the main k-mer) while also meeting the lowest bound of such motifs.

This defined the 6-mer groups within two mismatches. Since a large number of 6-mers spectrum is created whose search with two mismatches across the sequences becomes a computationally intensive and time consuming step, a FM-Indexing and Burrows Wheeler Transformation (BWT) based inexact search step was applied. Since, parallelism through multiprocessing was also implemented, the search becomes more faster with available cores of CPUs.

4,096 possible combinations of 6-mers were individually searched in the peak data for every RBP. To select the most abundant 6-mers, the first criteria was its occurrence across at least 70% of the CLIP-seq peak region data. All the 6-mers which were occurring in at least 70% data were evaluated for their significance occurrence at p-value<=0.01 using binomial test. Many most frequently occurring 6-mers were found whose numbers varied for RBPs (from RBM39 (3) to ELAVL1(17)). The found significant spots for 6-mers for any given type worked as the seed which were subjected to bi-directional expansion. This expansion step every time evalauted the similarity between the expanded region and checked for minimum similarity cut-off of 70% across the considered seed regions which were expanding. The 6-mer seeds were expanded at every found position until they were satisfying both the criteria. The final step resulted into the most frequently occurring elongated k-mers with maximum possible elongation with both criteria met. After elongation, the best scoring expanded k-mer family for each RBP was considered as the primary motif in the RNA sequences interacting with the given RBP. It was found that at least one such primary motif existed for all the RBPs considered in this study, barring four RBPs. These primary motifs were occurring in at least in 70% of the data with high significance. The size of primary motifs varied from 6 bases to 10 bases for different RBPs. The most abundant motif was based on UCUGCAG for ALKBH5 (92.27%), where as the least abundant motif was based on CCUGGAGG for SLBP protein (Supplementary Data 1 Sheet 3).

There were four different RBPs *viz.* FXR1, SND1, ILF3, and U2AF1 which did not have any single seed k-mer occurring in at least 70% of the data. This suggested the possibility for multiple motifs working in mutually exclusive manner. It was found that two different motif groups for these four RBPs were working almost in mutually exclusiveness manner with small fractions of overlaps in their instances. The overlap levels between these two motif groups’ instances were: FXR1 (7.8%), SND1 (6.8%), ILF3(8.25%) and U2AF1 (9.5%) instances (Supplementary Data 1 Sheet 4). However, for these cases the found 6-mers could not be expanded further as at least 10% of the data was lost due to this.

This way, the most significant motifs present in the cross-linking data of all these RBPs were discovered which could act as anchor in contextual form. It was interesting to observe that the identified motifs could be clustered into various groups based on their similarity. For every RBP, the motifs obtained from their respective sequences were used to develop their position weight matrices and logos which were compared with each other for similarity based clustering. This resulted into 28 clusters of RBPs where RBPs belonging to same cluster shared good level of similarity for their prime motifs (Supplementary Figure 1). Such display of grouping among RBPs is reflection of unity in diversity phenomenon as well as strongly suggest that how much of importance contextual factors could be for RBP-RNA interactions that despite of sharing similarity in their main motifs the binding appeared highly contextual. This also transpires from the study done on the flanking regions of these main motifs for the RBPs belonging to the same cluster. The di-nucleotide, pentamers and heptamers based information content strongly varied among themselves for many cases. The upcoming sections will present some related information on this.

When these motifs were mapped back to the genome in order to derive the contextual information, several of them hinted for coexistence of secondary supporting motifs for any given RBP. Such cases were studied further for co-occurrence of motifs where the most dominant motif would be supported by some other predominant secondary motif. All those sequences where the dominant motif existed were also searched for the supporting secondary motifs. Obtained co-occurring motif pairs were further evaluated to measure the similarity between them using Jaccard similarity index based approach. The method utilizes the position weight matrices of co-occurring motifs for alignment considering relative shifts to recognize similarity between two motifs (40). All co-occurring motif pairs possessed similarity score < 0.2 ensuring different motif partners being evaluated instead of same motif repeating itself. Sequence regions where the motifs co-occurred displayed high statistical significance of co-occurrence rate for the motif pairs for any given distance(p-value<<0.05; KS-test). For all RBP models, big difference was observed for the distribution of co-occurring motif pairs when compared to the random sequence regions, strongly supporting the existence of co-occurrence of motifs in RBP binding models of RNAs. Figure 2 illustrates some of these cases. In this way, 178 statistically significant co-occurring motifs pairs out of 297 motif pairs for 127 RBPs were obtained, strongly suggesting again that context holds importance in RBP-RNA interactions. Co-occuring motif details for the RBPs is given in the Supplementary data 1 Sheet 8. Further these motifs were also checked for frequency ratio >1 as discussed in method section. All 178 statistically significant co-occurring motif pairs were found to have frequency ratio (FR) > 1. These co-occuring motifs were analyzed for the region flanking 75 bases from both sides of the prime motif. The reasons for considering this region becomes more clear in the following next section.

**Figure 2:**
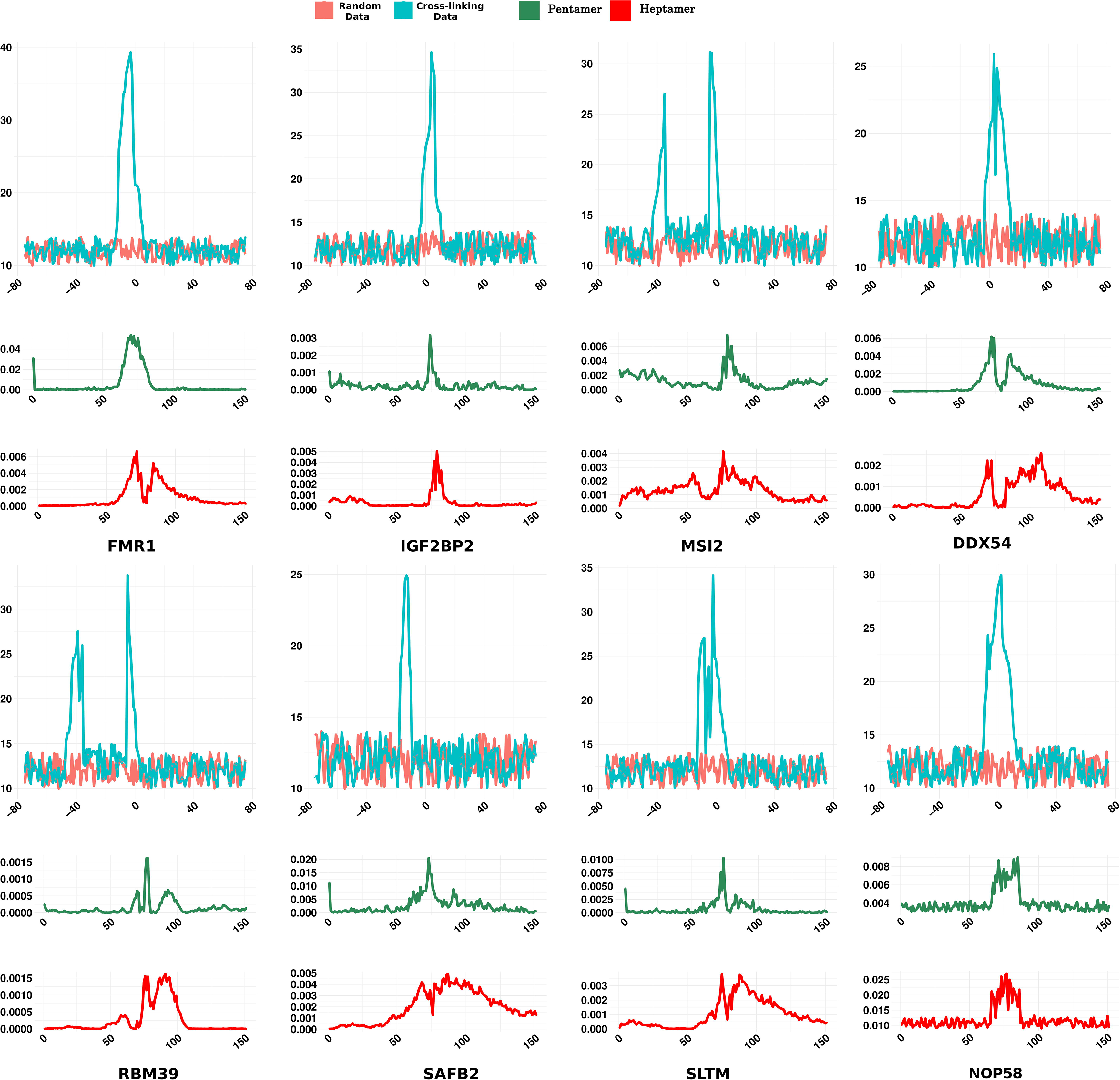
The co-occurring motifs positional preference. The plots are showing the position specific existence of the co-occurring motifs with respect to the prime motif (coordinated at “0”). F-score values of other contextual features like position specific pentamers and heptamers distribution reflect this to some extent. Most of these RBPs exhibited some secondary motif which co-occured with the prime motif in a position specific manner.

The motifs reported in the present study were compared with the experimentally reported motifs. Most of the motifs found in this study matched with the experimentally reported motifs. However, it was also observed that several of these experimentally reported motifs were not the prime motif reported here but matched to other lower ranked motifs which either co-occured with the prime motifs or were exclusively present, covering comparatively lesser amount of CLIP-seq data than the prime motifs reported in the present study. Their occurrence in the cross-linking data varied from 34.07% to 81.32% while the prime motifs reported in this study mostly covered at least 70% of CLIP-seq data. Figure 3 provides a snapshot of the comparison between experimentally reported motifs with motifs identified in the present study.

**Figure 3:**
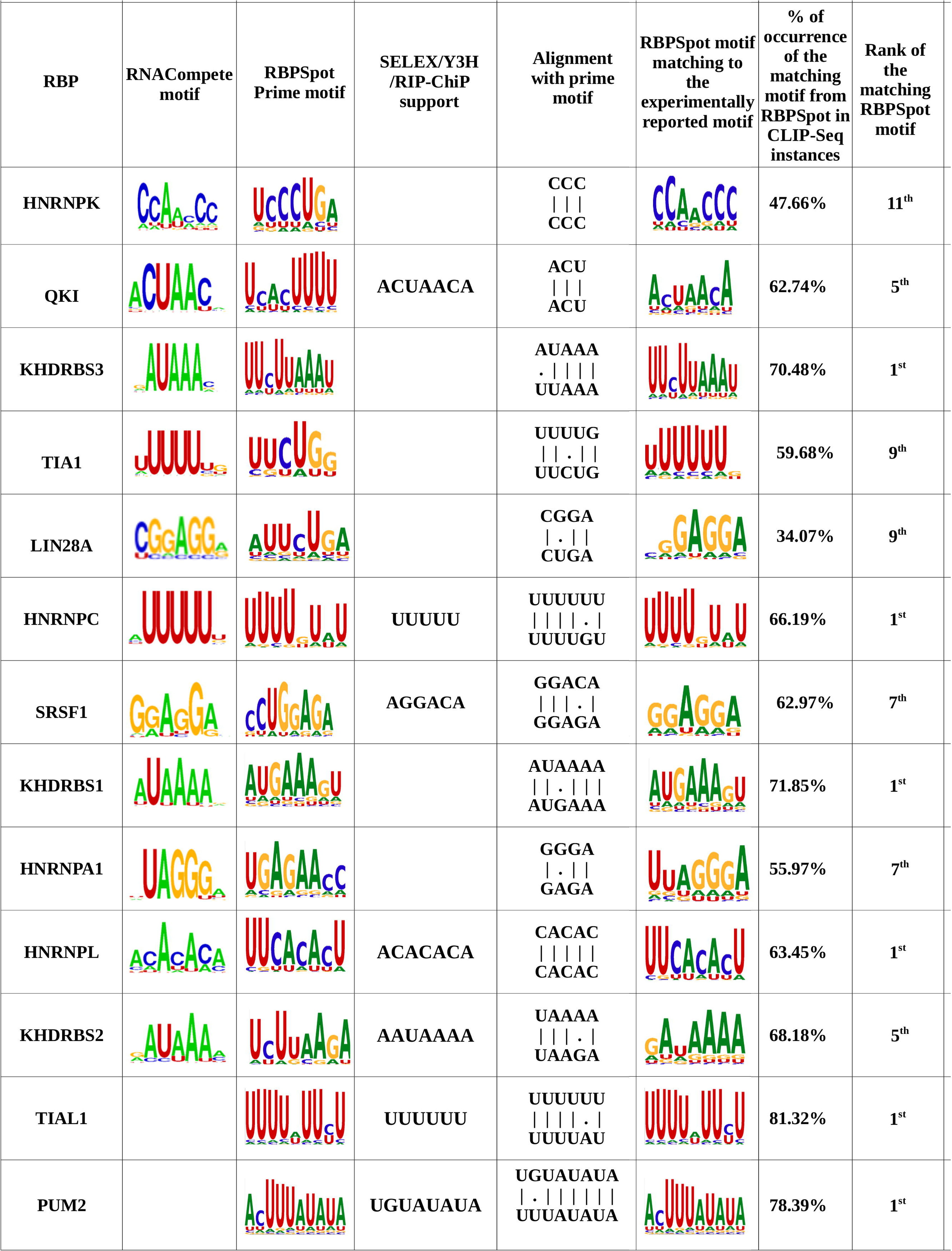
Comparison between experimentally reported motifs and motif identified in the present study. Most of the previously reported motifs for the RBPs were detected by the approach presented in the current study. However, it also observed that several of previously reported motifs are not the prime motifs but comparatively cover lesser CLIP-seq data than the prime motifs identified in the present study. The last three columns show the matching motifs similar to the previously reported motifs, their status in CLIP-seq data coverage, and the corresponding motif rank.

### Consideration of expression data for targets helps in building more realistic data-sets

The discovered motifs above work as a point to zero upon to consider the potential significant interaction spots in the RNA. However, such motifs alone can’t hold much higher stake than that as they may appear in the non-binding regions also, though found statistically significant for the binding regions. Evaluation of their context for their functional role thus becomes essential. In this regard, 75 bases from both the flanking regions were considered where the motif region worked as the anchor. Previously, it has been found that ∼75 bases of flanking regions around the potential interaction sites in RNAs capture the local environment for structural and contributory information effectively (20). Also, RBPs which interact with the RNA through multiple domains use multiple interaction sites which are usually concentrated around a local region instead of being long distanced interaction spots. Thus, uniform length sequences with flanking regions were obtained for every individual RBPs which varied for different RBPs depending upon the length of their anchor motifs. This also led to the construction of positive and negative instances datasets, simultaneously. The number of positive instances differed for the RBPs depending upon their available cross- linking sequencing data, ranging from 1,309 (EIF4A1) to 8,48,680 instances (AGO1-4). This covered a total of 19,547 genes experimentally confirmed as targets of these RBPs. Total number of instances was greater than total number of peak data for most of the RBPs due to multiple occurrence of motifs on a single sequence.

Identifying suitable negative dataset candidates becomes a more crucial task. And it is where most of the previously developed tools have gone too soft and mostly ended up selecting random sequences, which actually does not help to divulge more information. As transpires from above discussions and results, there are many spots across the transcriptomes which posses sequences similar to the interaction motifs but they yet not interact. In usual, chances of finding shorter motif themselves is higher in the random data. In such scenario, considering random sequences really does not add significantly to the purpose of discrimination and does not answer the question raised above. In order to build a better negative dataset, it is better to pick those candidates as negative instances where the region similar to the main motif is present and creates a strong confusion matrices to build a more natural and robust model. Therefore, to create the negative set for RBPs two different kind of strategies were used. In the first strategy we used RNA-seq data for the same condition for which we had the cross-linking data available for the given RBP. Those RNAs were selected which were expressing themselves in the same condition but did not bind to the considered RBP and did not reflect in the CLIP-seq data. They were searched for the prime motifs of the RBP similar to the positive data cases and in similar manner 75 bases flanks were considered along with capturing the contextual information with more discrimination power. For the RBPs for which the negative datasets were created using this strategy are called Set A RBP datasets throughout this study. This way, the negative datasets for 74 RBPs were created (Supplementary Data 1 Sheet 5).

In the second strategy, the negative datasets were created for those RBPs which did not have similar condition RNA-seq data available for the considered CLIP-seq conditions. In such scenario, therefore, here those RNAs were considered which exhibited binding to their respective RBPs but they also had the motifs on other positions which did reflect in the CLIP-seq data and were also far away from such cross-linking regions. The logic behind is that such RNA sequences whose some regions exhibited binding to RBPs in CLIP-seq data make clear positive instances out of these regions as well as hold a simultaneous evidence that these RNAs were expressed in the given condition. Regions which display the interaction motif in these expressed RNAs but don’t bind to the RBPs become an apt case for negative instance consideration with high potential for contextual information unlike the usual random sequences. This particular set of negative dataset instances were called Set B. In this way, the Set B negative dataset were created for the remaining 57 RBPs (Supplementary Data 1 Sheet 6)). Rest of the analysis were same on both the sets of RBPs. This all also reinforces the view that any successful RBP-RNA interaction discovery approach can not be founded solely upon the motifs consideration but needs correctly designed context information extraction approach also which can be provided only after a better a negative instances consideration.

### Contextual information surrounding the anchored motif is critical for RBP binding sites recognition

Motif discovery and anchoring helped in selecting the more appropriate positive and negative instances from which contextual information and features might be derived. The contextual information came in the form of other co-occurring motifs, sequence specific information, position specific information, and structural/shape information which could exhibit sharp discrimination between negative and positive instances. Contextual information were derived from the features based on four major properties: (1) 7-mers frequency probability for each position, (2) 5-mers frequency probability for each position, (3) di-nucleotide densities in the region, and (3) Structural triplet frequency covering 27 combinations of structure triplets arising from the dot-bracket structural representation from RNAfold predicted RNA structures. Consideration of heptamer was for picking up any further sequence specific signals in the flanking region, where similar approach of inexact search was applied with at least 70% similarity match as was done for the prime motifs’ 6-mer seeds. Pentamers application was motivated from the recent findings which reported that pentamers capture the DNA shape very accurately (24). The nucleic acids shape has been found critical in the interactions with regulatory proteins which scan these shapes for their stationing. So far, this approach has been applied on DNA but hardly on RNAs. The DeepBind work had observed about the importance of using such kind of information which could be beneficial in future developments for the tools reporting RBP-RNA interactions (14). The dinucleotide densities have been found to be highly useful in indirectly evaluating the RNA structure and accessibility (20, 21). In fact, it has been found more promising than *ab-initio* RNA structure prediction. *Ab- initio* methods’ accuracy drastically falls with the length of RNA, and they are suitable for only short RNA sequences (9, 20). Pentamer and di-nucleotide frequencies capture better structural and shape information through base stacking and neighborhood contribution. Similarly, RNA structure triplet has been used widely in deriving the structural information of RNA for their propensity towards interaction factors, especially for miRNA:RNA interactions (41).

Various features generated based on the above mentioned properties were evaluated for their discrimination potential between the positive and negative instances. The most important top 100 features are given in supplementary data 1 sheet 9. Among them, the features originating from the dinculeotide densities appeared the most. Some pentamer and heptamer features were also present among these top features. Dinucleotide density reflects the structural and accessibility properties of the nucleic acids, as mentioned above. A very striking observation was also made here. Most of the positive instances flanking regions displayed enrichment of CG. Approximately 69% of RBPs target regions exhibited CG among the most dominant feature for each position. Where as for rest of the RBPs had UU and UA among the most prominent features. Besides this, it was also observed that RBPs which shared high similarity for their binding motifs and were clustered among the same group (Supplementary Figure 1) differed substantially for this contextual information and their flanking regions displayed different distribution patterns. Figure 4 presents an example of one such group, RBPs belonging to AGO4 cluster (Cluster 1). As can be noted in this figure also, CG is remarkably enriched for the binding site regions. Therefore, despite of having binding sites motifs they differ in their binding which is influenced by context. Also, the universal prominence of CG in the RBP binding regions reinforces the theory which suggests their regulatory roles in stationing the binding factors and supporting the binding motifs (42). Also, they may be studied further for RNA modification which are considered critical for RBP binding dynamics.

**Figure 4:**
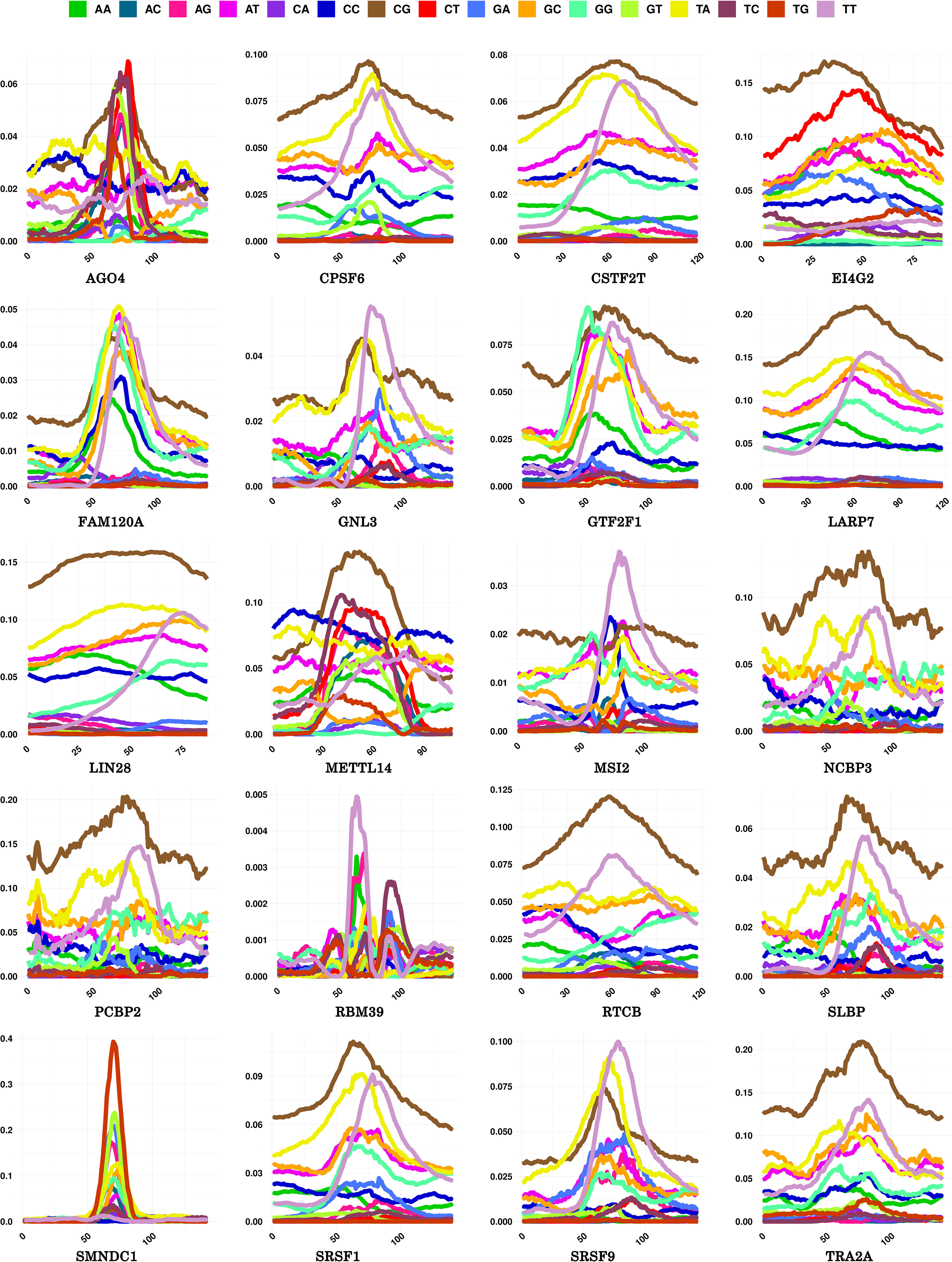
F-score distribution of dinucleotide densities at different positional windows for the target regions and their flanks for Cluster#1 members. Context specific dinucleotide density distribution emerged among the most important features for all RBPs taken in this study. Their densities worked as important features at variable windows and distances for different RBPs. Here, Cluster# 1 members data is shown. They shared high similarity among themselves for their prime binding motifs, yet their contextual information and density profiles differed a lot. Enriched contextual “CG” distribution of these regions was found consistently distinguished property for the regions binding the RBPs.

For 12 RBPs, pentamers were also found in the top 20 features for different positions whereas heptamers were found for 10 RBPs in the top 20 features. Among top 100 features, almost in 90% cases heptamers and pentamers marked their presence. Significant difference was observed between the positive and negative instances with respect to the F-score for positions which also suggest that substantial amount of information is being held by the flanking regions around the binding motif, which may be one of the determinant for contextual interactions between RBP and RNA. A series of t-tests between the positive and negative instances for various features also supported this. Biologically, heptamers and pentamers were expected to reflect any supporting co- occurring motifs near the prime anchored motif. Pentamers, specifically, were considered to capture the shape properties, which too have been called important in protein and nucleic acids interactions, more so in cases where sequence motifs are not clear or prime (14, 24). A closer look with these pentamers and heptamers revealed that for many RBPs binding sites, they were prominent in the flanking regions where the co-occuring secondary motifs existed (Figure 2). Though heptamers were found more reflective to this phenomenon. As could be expected now, these information properties from the flanking regions looked highly promising for identification of a true binding site. The impact of each of these properties on discrimination capacity between true binding sites and negative sites was also clear when evaluated directly on the machine learning models for performance, as transpires in the following section.

### DNN implementation of the RBP binding site models consistently achieved high accuracy

Before combining the features to build the collective models for RBP-RNA interactions, one more assessment of contributions by the above mentioned properties in discrimination was done. Classification assessment was made for each given properties separately before joining them together while using XGBoosting. This was done to get the preliminary idea about the individual contribution made by each of the contextual properties towards the accurate classification and how important they looked in the process of accurate recognition of the binding spots. For the pentamers based classification the accuracy varied from 60.23% (U2AF2) to 82.01% (FKBP4) for Set A RBPs with an average of 69.8% accuracy. For heptamers it varied from 65.01%(FXR1) to 88.72% (FXR2) with an average accuracy of 76.49% for set A RBPs. Similarly, for set B RBPs pentamer accuracy varied from 55.6% (DHX9) to 86% (EIF4A1) with an average of 66.7% accuracy. For heptamers it varied from 57.23% (MOV10) to 97.47% (EIF4A1) with an average accuracy of 77% for Set B RBPs. For structure triplets we used different window size but none of the windows achieved more than 63.39% accuracy for any RBP, clearly supporting our above made observation that *ab-initio* structure prediction derived features don’t add much value due to their innate limitations. Therefore, this feature was not further taken for the final model building. Accuracy for di-nucleotide densities based classification varied from 63.04% (FMR1) at 43 window size to 88.6% (RBFOX2) at 71 window size with an average of 75% accuracy at different window sizes which varied from 17 to 103 for Set A RBPs. Similarly, for set B RBPs the accuracy of di-nucleotide density based classification varied from 61.46% (DHX9) at 71 window size to 90.57% (EIF4A1) at 91 window size with an average of 75.25% accuracy (Supplementary Data 1 Sheet 5,6). The results here displayed concordance with the observation made in the previous section where importance of contextual dinucleotide density information based features emerged as the most important ones for the binding sites detection. Figure 5(A) presents the violin plots for the accuracy distributions observed for the classifications done by each of these properties for all the RBPs.

**Figure 5:**
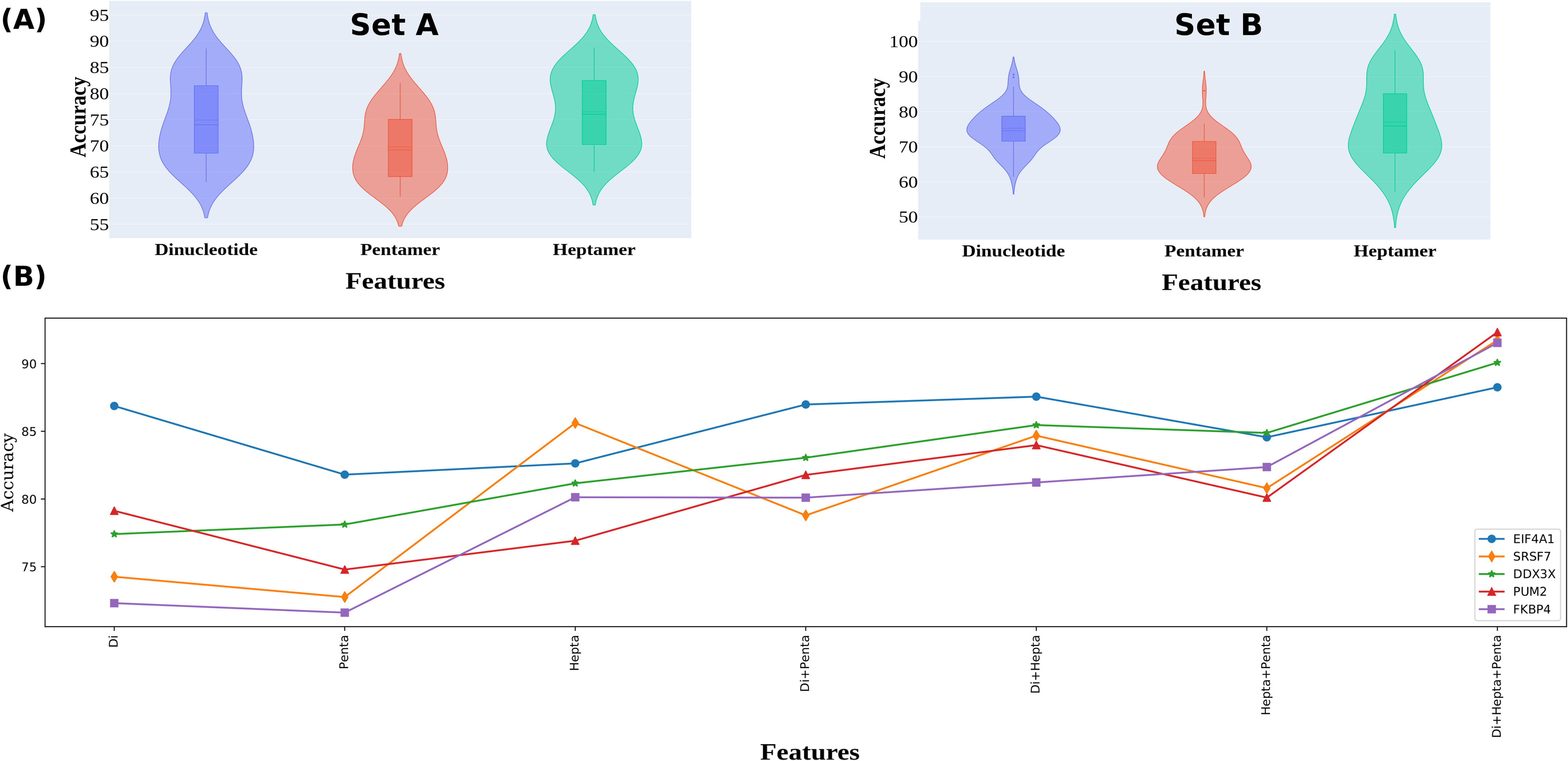
Assessment for three main properties in discriminating between the negative and positive instance,(A) Violin plot distribution of accuracy when dinucleotide, pentamer and heptamer were used alone for set A and set B RBP. (B) Impact of combination of the dinculetodie, pentamers, and heptamers properties based features. These features appeared highly additive, complementary to each other as the performance in accurately identifying the binding regions increases substantially as these are combined.

With this all, it was pretty evident that the selected properties and their features had strong discriminatory strength, barring the RNA structural information derived through *ab-initio* structure prediction method. All the features originating from these qualifying properties were combined together to build the final models of RBP-RNA interaction targets.

After getting optimum window size for di-nucleotide densities, we combined these three features (pentamers probabilities, heptamers probabilities and di-nucleotide densities) together to build the final models. The final models were built using Xgboosting as well as DNN. The reason for considering these two different approaches are that: 1) they reflect two different learning approaches: Shallow and Deep, 2) They complement each other as Xgboost works good for the cases with comparatively lower training data while DNN performance is good where learning data is higher, 3) Both the approaches work very good for conditions where the dimensions are high, as was with this study.

Combining of the features based on above mentioned properties was done in a gradual manner in order to see the additive effect of them on the classification performance. As it is apparent from Figure 5(B), which showcases the DNN classifier’s performance for five RBPs, the performance of the classifiers kept increasing on the addition of more features, where consistency also increased as can be seen through the band width of the plots for the five RBPs. Here also, the dinucleotides based contextual features emerged most critical as the biggest leap in the performance was noted when it joined the heptamers and pentamers based features. Any pair of these three properties features gave almost similar performance, but sharpest rise was observed in the performance when contextual dinucleotide information based features were added to the pentameric and heptameric features.

After combining the features we had 1,198 (ZNF184) to 2,544 (EIF4A3, EIF4G1, EWSR1,HNRNPD, HNRNPL, KHDRBS3, NOP58 etc.) features for individual RBPs. The feature numbers varied due to different sized best performing dinucleotide densities windows. These features were used in Xgboost machine learning where the average accuracy of 85.07% (Avg. AUC: 85.06%, Avg F1-Score: 84.64% Avg MCC:79.26) was obtained and where the values varied from 79.19% (FXR2, AUC: 79.19%, F1-Score: 78.58% MCC:66.38) to 90.81% (RBM47, AUC: 90.80%, F1-Score:90.49 % MCC:83.17)) for Set A RBPs. It was found that the average accuracy of 84.08% (Avg. AUC: 84.07%, Avg F1-Score:82.66 % Avg MCC: 69.07) was obtained for Set B RBPs, where accuracy values varied from 66.34% (MOV10, AUC: 66.34%, F1-Score: 64.74% MCC:42.58) to 96.78% (EIF4A1,AUC: 96.48%, F1-Score: 96.40% MCC: 92.37). The same set of the combined features was also used in the DNN implementation. DNN works better with higher dimensions and instances to learn from. In the input layer combined features were used where as two hidden layers gave best performance and the number of nodes per hidden layer varied from 700 to 1,300. Details of implementation are already given in the methods section. DNN achieved an average accuracy of 92.25% (Avg. AUC: 92.64%, Avg F1-Score: 91.97%, MCC:84.52%) for Set A RBPs which was much higher than XGBoost. Whereas for Set B RBPs an average of 83.47% (Avg. AUC: 89.61%, Avg F1-Score: 83.18%, Avg MCC:67.34%) accuracy was achieved by the DNN models, which was slightly lower than XGBoost. Complete performance details can be found elsewhere (Supplementary Data 1 Sheet 5,6).

In general, it was apparent that DNN approach was sensitive towards the volume of training instances as it was found performing better where number of instances were higher. But the biggest impact on performance was observed was for the granularity of dataset creation. Performance of DNN was specially more marked here, as can be seen from its performance plot on Set A datasets. On Set A, the DNN models performance hardly touched below 90% accuracy. Even XGBoost’s performance was better with Set A when compared to Set B. It needs to be recalled that Set B was made for those RBPs for which the RNA-seq data was not available for the considered CLIP-seq conditions. In such scenario, those RNA were considered to generate the negative instances whose some regions were present in the CLIP-seq data suggesting their expression. From the same RNA, those regions were selected which were having the prime motifs but yet not binding to the RBP and not reflected in the CLIP-seq data and were distant from such binding regions. While Set A negative instances were clearly those regions which were expressed during the CLIP-seq experimental condition and possessed the prime motif but no region of the RNA itself bound to the RBP. Thus, though the over all performance with Set B was still good and better than the datasets used by the compared tools as transpires in the next section, it same time reflects that how important it is to have a refined data-set like Set A. This is possible that some instances covered as negative instances in Set B could be contributing to the RBP-RNA interactions or could not be captured in the CLIP- seq experiments. Yet, as transpires from the various performance metrics plots across various RBPs given in Figure 6 and AUC/ROC plots given in Figure 7, the developed approach in this study, named as RBPSpot, showcases a consistently high and reliable performance for a large number of RBPs. It also provides the largest number of models for RBPs binding developed from CLIP-seq data to this date.

**Figure 6:**
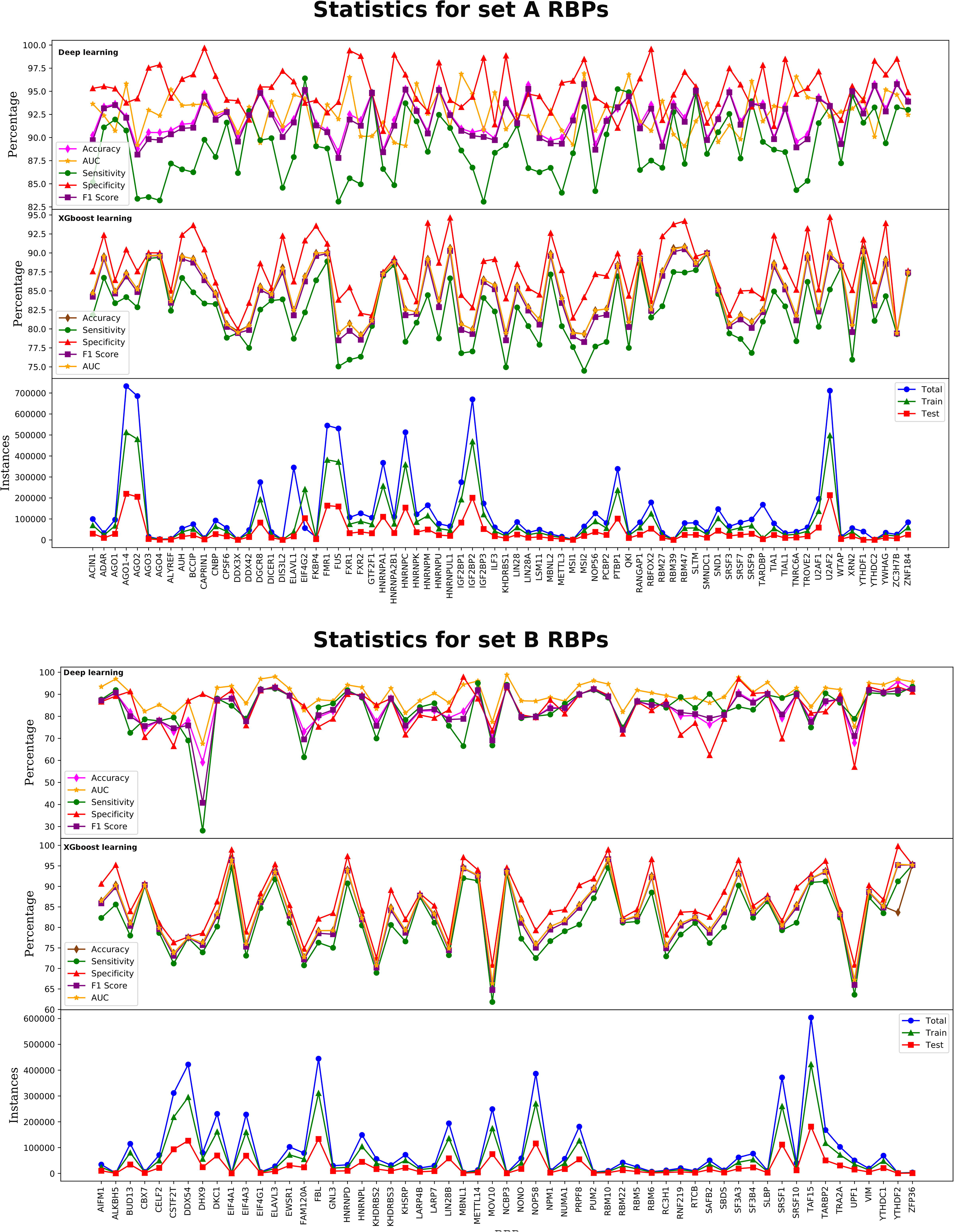
Performance metrics for RBPSpot. (A) First plot showing the accuracy, AUC, sensitivity, specificity, F1 score for the DNN model for set A RBPs. The second plot is showing the same metrics for the gradient boosting method. The third plot is showing the corresponding instances in the test, train and in total data for set A RBPs, (B) The first plot is showing the accuracy, AUC, sensitivity, specificity, F1 score for the deep learning models for Set B RBPs, where the second plot is showing the same metrics values for the gradient boosting method with Set B RBPs. The third plot is showing the number of instances in the test, train and in total data for Set B RBPs. RBPSpot scored highly on all the performance metrics where the most remarkable thing about it was its consistent performance across a large number of RBPs and dataset.

**Figure 7:**
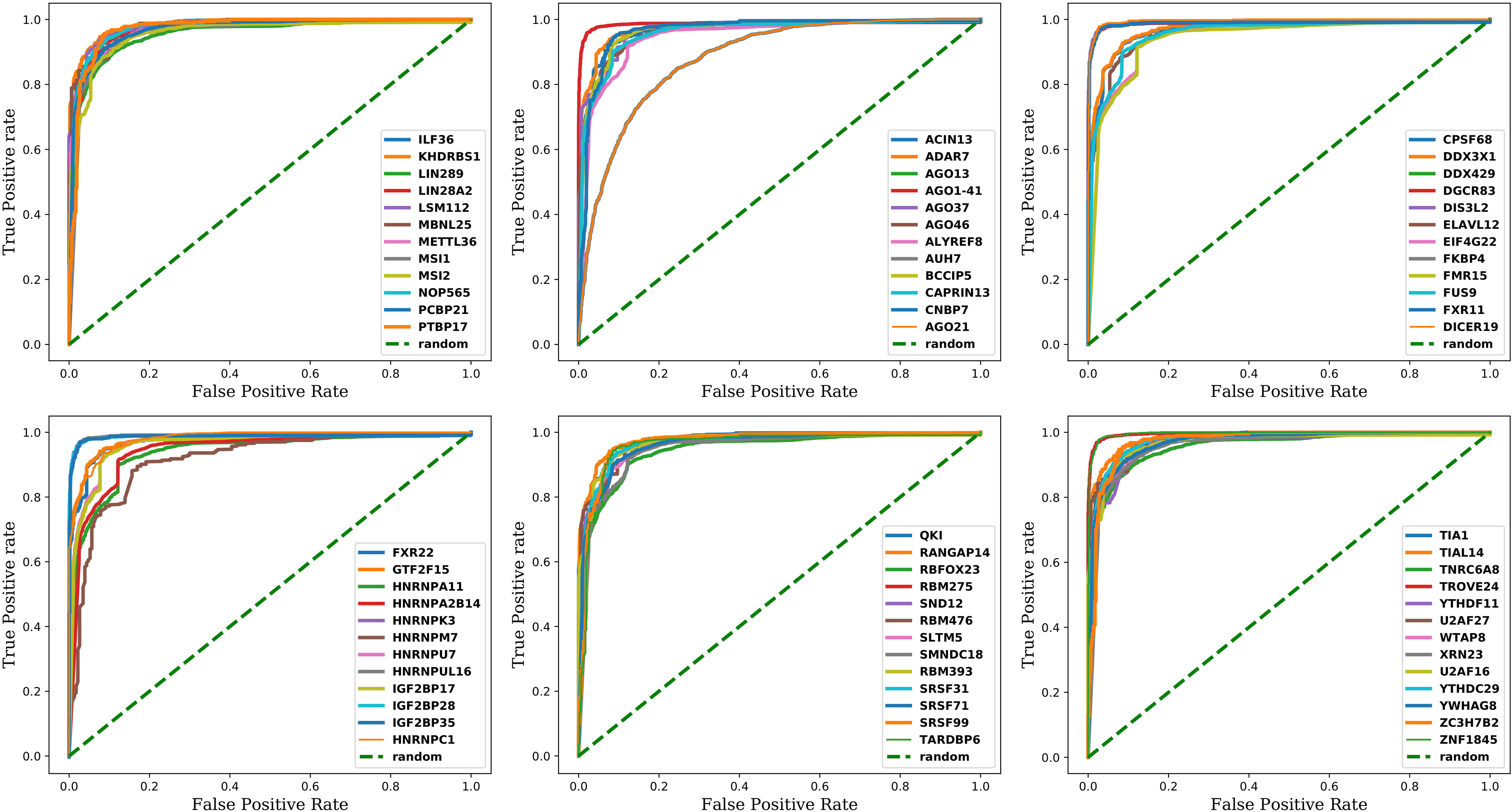
AUC/ROC plots for Set A RBPs. The AUC/ROC plots for the deep-learning models for some of the RBPs clearly showcase the robustness and highly reliable performance of the implemented DNN models.

### Comparative benchmarking: RBPSpot consistently outperforms all the compared tools

A very comprehensive benchmarking study was performed where RBPSpot was compared with five different tools, representing different approaches of RBP RNA interaction detection: RBPmap (probabilistic approach), beRBP (Random Forest machine learning bases claiming highest accuracy in its category), DeepBind (the first deep-learning based approach), iDeepE and DeepCLIP (representing some very recent and more complex deep-learning based tools). Besides this, the benchmarking has also considered three different datasets as this work also presents a new dataset while underlining the importance of better datasets in creating better models as well as to carry out a totally unbiased assessment of performance of these tools on different datasets.

Thus, the first dataset considered in the benchmarking study was derived from the RBPSpot dataset. Only those RBPs were considered for comparison for which at least one tool had model built, besides RBPSpot itself. This way comparison was done for 52 RBPs. The second dataset considered was the one evolved during development of Graphprot software which has been used largely by various other datasets for model building and performance benchmarking purposes. The third dataset used in this benchmarking study was the one used by beRBP software which too has been used by many other tools for the same purpose. Details about these datasets have already been discussed above and in the methods sections.

All these six software were tested across all these three datasets and RBPSpot outperformed all of them across all the datasets, and for all the performance metrics considered (Figure 8). Figure 8 gives a detailed view of the data analysis of this benchmarking across the three datasets studied for all these software. RBPSpot scored the average accuracy of 88.43% and the average MCC value of 0.77 on RBPSpot dataset, the average accuracy of 91.63% and the average MCC value of 0.83 on Graphprot dataset, and the average accuracy of 88.9% and the average MCC value of 0.74 on beRBP dataset. Among all the considered performance metrics, MCC stands as the most important one as it gives high score only when a software scores high on all the four performance parameters (true positive, false positive, true negative, false negative). A good MCC score signifies the robustness of the model and its performance consistency. RBPSpot emerged as the most robust algorithm among all these compared software with very high consistency of performance. As it is visible from the score distribution for all the metrics, RBPSpot also exhibited least dispersion of scores for all the studies RBPs and for all the three datasets, confirming the precise performance achieved by RBPSpot compared to other tools.

**Figure 8:**
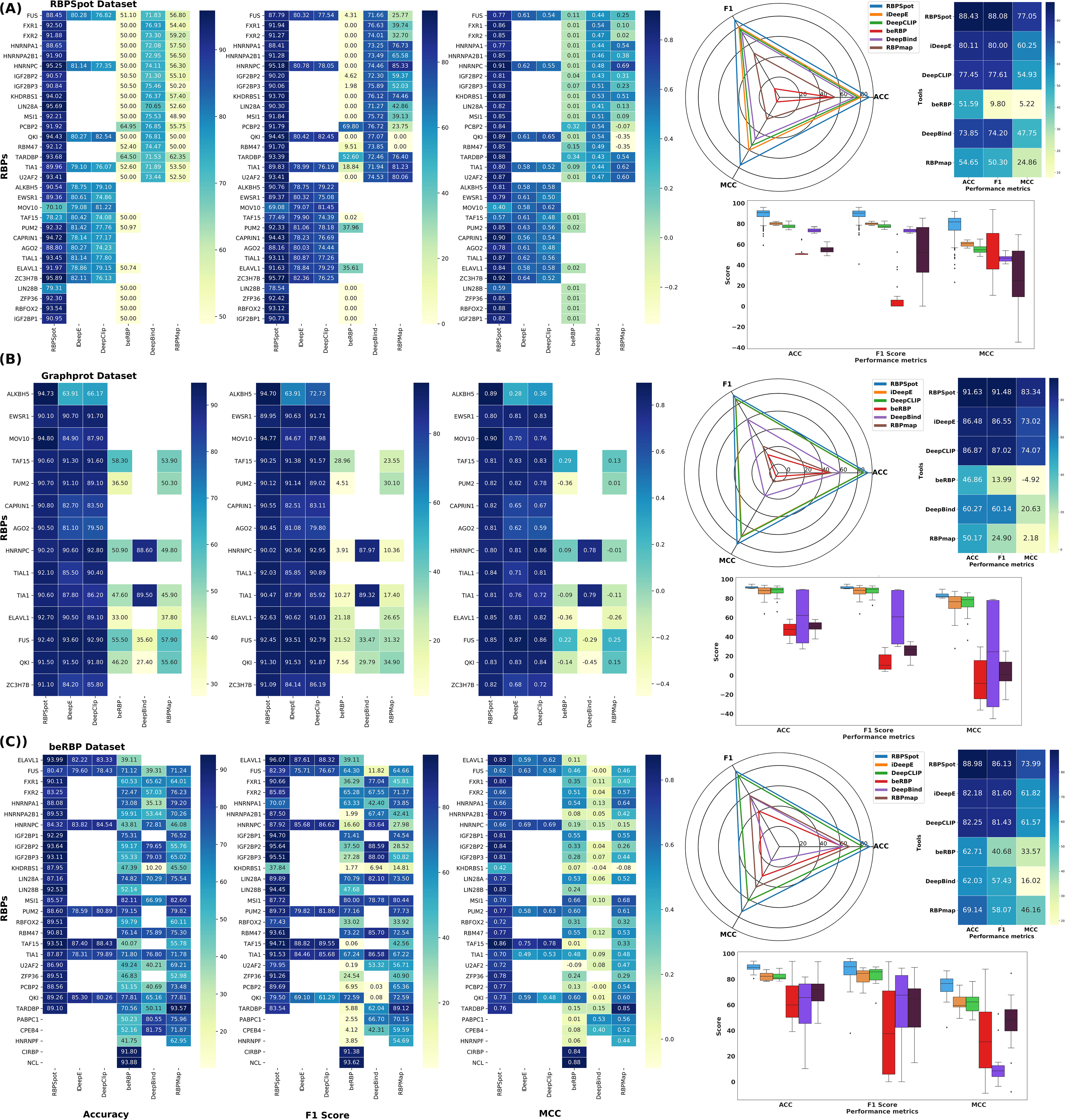
Comparative bench-marking results of RBPSpot when compared to beRBP, DeepBind, RBPmap, iDeepE, and DeepCLIP for three different datasets. (A) Bechmarking result on RBPSpot dataset, (B) Graphprot dataset, and (C) beRBP dataset. Each these datasets performances was evaluated for various performance metrics where the heatmaps are for accuracy, F-1 score, and MCC values for each dataset for some of the evaluated RBPs. The rightmost plots are radar charts view of the average Accuracy, F1 score, and MCC attained by each software for the corresponding dataset. The last plot is the box plot which provides the average distribution of these metrics scores. From the plots it is clearly visible that for all these datasets and for almost all of the RBPs, RBPspot consistently outperformed the compared tools for all the metrics. More all the radar plots it scored the highest and nearest to the isosceles triangle suggesting consistent and better average performance also. The box plot suggests that RBPSpot not only performed best in overall but also the dispersion of its various metric scores were much lesser than other compared tools. Some of these tools exhibited enormous variation in the distribution of their metric score suggesting unstable performance by them.

After RBPSpot, the best performance was observed for the complex deep-learning based software iDeepE and DeepCLIP. On RBPSpot dataset, iDeepE performed better than DeepCLIP, but for other two datasets they attained almost similar metrics scores for performance. Undeniably, they emerged far superior than their deep-learning predecessor, DeepBind, and other compared tools. They even displayed much smaller dispersion of their scores than other compared tools. However, RBPSpot’s performance points out that more appropriate features may be learned through training on biologically relevant properties to derive better discrimination power using machine learning approach, which can be amalgamated with Deep Neural Nets with much lesser complexity and superior performance than applying complex deep-learning layers to automate feature extraction. The observations made in the introduction part of this work appeared true in this study that such complex deep-learning approaches score good on unstructured data where clear features identification and extraction is difficult to be done by expert and automation is required for feature extraction. The problems where features are identifiable and can be structured, simpler machine learning models may outperform the complex deep-learning approaches.

The above mentioned benchmarking was done for all the tools while keeping their original training dataset and models for RBPs. Most of the existing tools don’t provide the option to build user specified models of RBPs using their algorithms but come with their own pre-built models. This limits the scope to test the algorithms with different combinations of datasets. Fortunately, the two best performing tools after RBPSpot, iDeepE and DeepCLIP, provided this scope where the users may build their new models with their own datasets. Also, since these two tools performance were next to RBPSpot, they stood as a natural choice to study the performance impact with datasets variations. Both iDeepE and DeepCLIP have implemented Graphprot dataset for their original model building. For this part of benchmarking study the training and testing datasets of RBPSpot, iDeepE, and DeepCLIP were swapped and studied for four different combinations of training and testing datasets: RBPSpot training and testing datasets, RBPSpot training and Graphprot testing dataset, Graphprot training and testing datasets, Graphprot training and RBPSpot testing dataset.

Figure 9 presents the results for this part of benchmarking where RBP models were rebuilt and tested using the four different combinations of training and testing datasets. RBPSpot outperformed the remaining two software, iDeepE and DeepCLIP for all the combinations of datasets, for all the considered performance metrics. Like the previous benchmarking study, here also RBPSpot scored the highest among all the software for all the combinations of datasets with a remarkable consistency. As transpires from the kernal density plots in Figure 9, RBPSpot maintained its least variability and dispersion of performance scores and continued to display it strong balance in detecting the positive and negative instances with high and similar level of precision. This was reflected by high scoring on all the four parameters of performance resulting into consistently highest MCC values, which confirmed the robustness of the algorithm. Also, it was observed that performance of all the compared software was better when RBPSpot dataset was used for training. The original implementation of iDeepE and DeepCLIP have used Graphprot dataset. Both these software performed better when their original dataset for model building was replaced by RBPSpot training dataset, underscoring better and more realistic composition of RBPSpot dataset.

**Figure 9:**
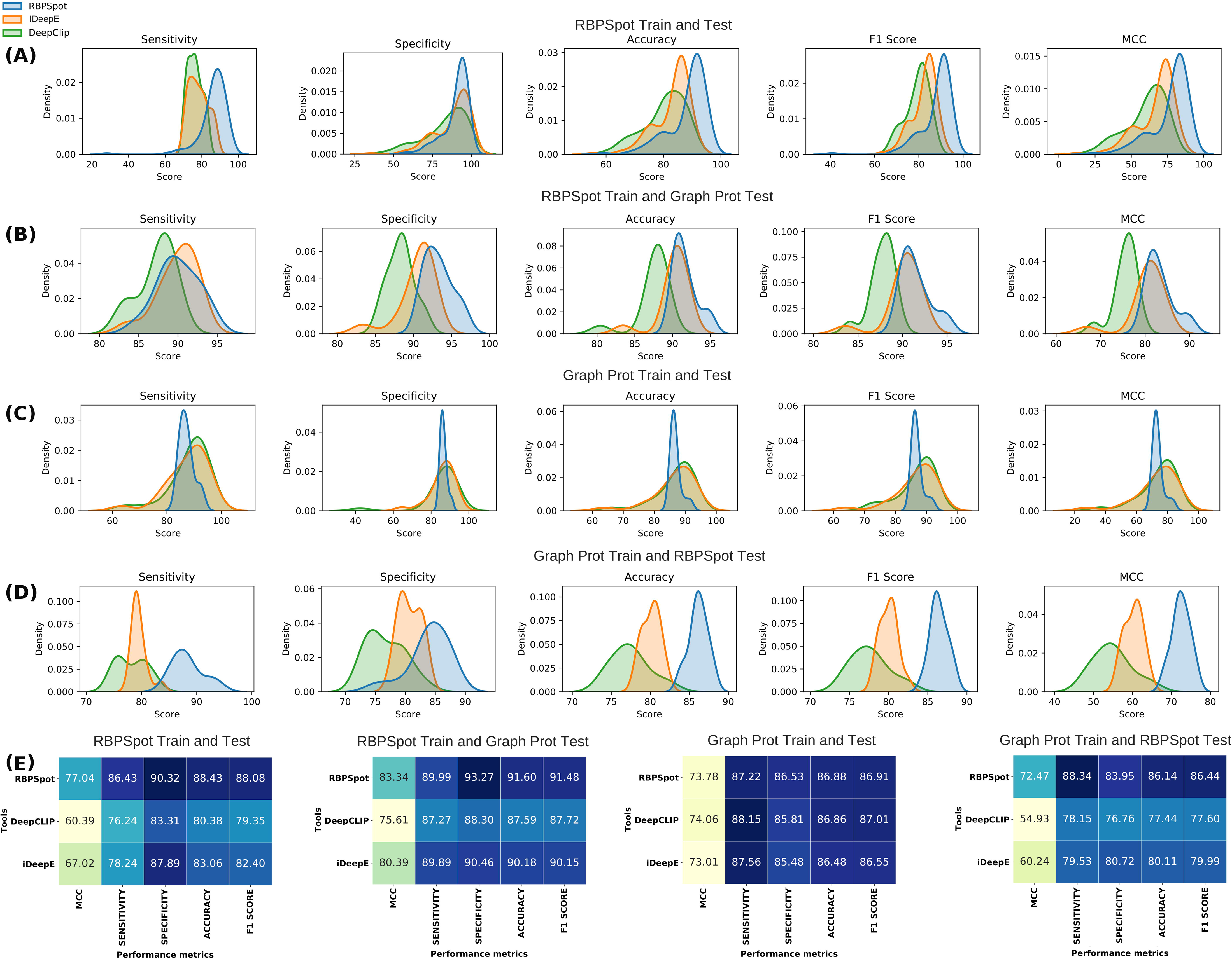
Performance benchmarking with different combinations of train and test datasets. In this part of performance benchmarking the impact of datasets was also evaluated. Since this part required rebuilding of RBP-RNA interaciton models from the scratch and from the provided user defined data, only two other tools other than RBPSpot qualified this criteria (DeepCLIP and iDEEPE). These tools provide the capability to build new models from user given datasets. These tools were originally developed on Graphport dataset. Therefore, in this part of benchmarking RBPSpot and Graphprot datasets were considered and 4 different train-test datasets combinations were studied. Distributions for various performance metrics for the compared tools and the corresponding datasets have been given as Kernel densitiy plots: (A) RBPSpot train and test, (B) RBPSpot train and graphprot test, (C) Graphprot train and test, and (D) Graphprot train and RBPSpot test. For every such combinations, the average performance metics scores are given in the form of heatmap (E). The plots clearly underline that RBPSpot consistently outperforms the two tools for all the metrics on all these different combinations of train and test datasets, where again the consistent and precise performance of RBPSpot was an important observation, Consistently high MCC scoring by RBPSpot underlined it as a robust and balanced algorithm where dispersion in performance metrics was least. Also, the performance of all the compared tools increased when RBPSpot dataset was used in training, clearly suggesting the importance of having a right dataset. RBPSpot dataset presented here emerged as a better dataset for such studies.

The benchmarking done here stands among one of the most comprehensive ones. It looked into various aspects of performances and has involved a large number of RBPs for comparison as well as evaluated the role of datasets in performance. RBPSpot consistently scored high across all the comparative tests and clearly outperformed the compared tools. The full details and data for the benchmarking studies are given in supplementary Data 1 Sheet 10-15.

### Structural and molecular dynamics analysis supports the RBP binding site models

Depending upon the availability of complete experimentally validated 3D structures in PDB database, structures for 13 RBPs (IGF2BP1, DIS3L2, CNBP, SRSF3, FKBP4, KHDRBS1, LIN28A, CAPRIN2, DICER1, GTF2F1, HNRNPC, CPSF6 and AGO2) were selected for the structural interaction analysis for the identified binding sites (43). In order to examine conformational variations of the RBPs within the hydrated controlled environment, the root-mean- square deviation (RMSD) of the atomic positions of RNAs containing motif with respect to RBP backbone were calculated and compared with the RNA complexes without the prime motif. In comparative analysis of RMSD measures these RBPs complexes were considered with three different RNA sequences for each RBP. These sequences were randomly selected from positive datasets having 75 bases flanking regions. To analyze the structural behavior of RBPs and their complexes, 20 ns simulation job was performed. For this purpose, selected RBPs and complexes were immersed in the cubic boxes of varying dimensions based on the system size. Prior to the energy minimization process, different charged molecules like NA^+^ or Cl^-^ were added to neutralize the system (44).

Once the simulation was finished, the last step was to analyze the simulation result in term of RMSD plot during the course of simulation for 20ns. RMS module in GROMACS was executed while choosing “Backbone” for least-squares fitting and “RNA_Heavy” for the RMSD calculation. By doing so, the overall rotation and translation of the protein was removed via fitting and the RMSD reported about how much the RNA position varied relative to the protein. This is considered as a good indicator of how well the binding pose was preserved during the simulation. Comparative analysis of RMSD trajectories of 13 different RBPs-RNA complexes with three replicates each for the two conditions clearly suggested that the presence of the identified prime motifs was giving stability to the RBP-RNA complexes (Figure 10).

**Figure 10:**
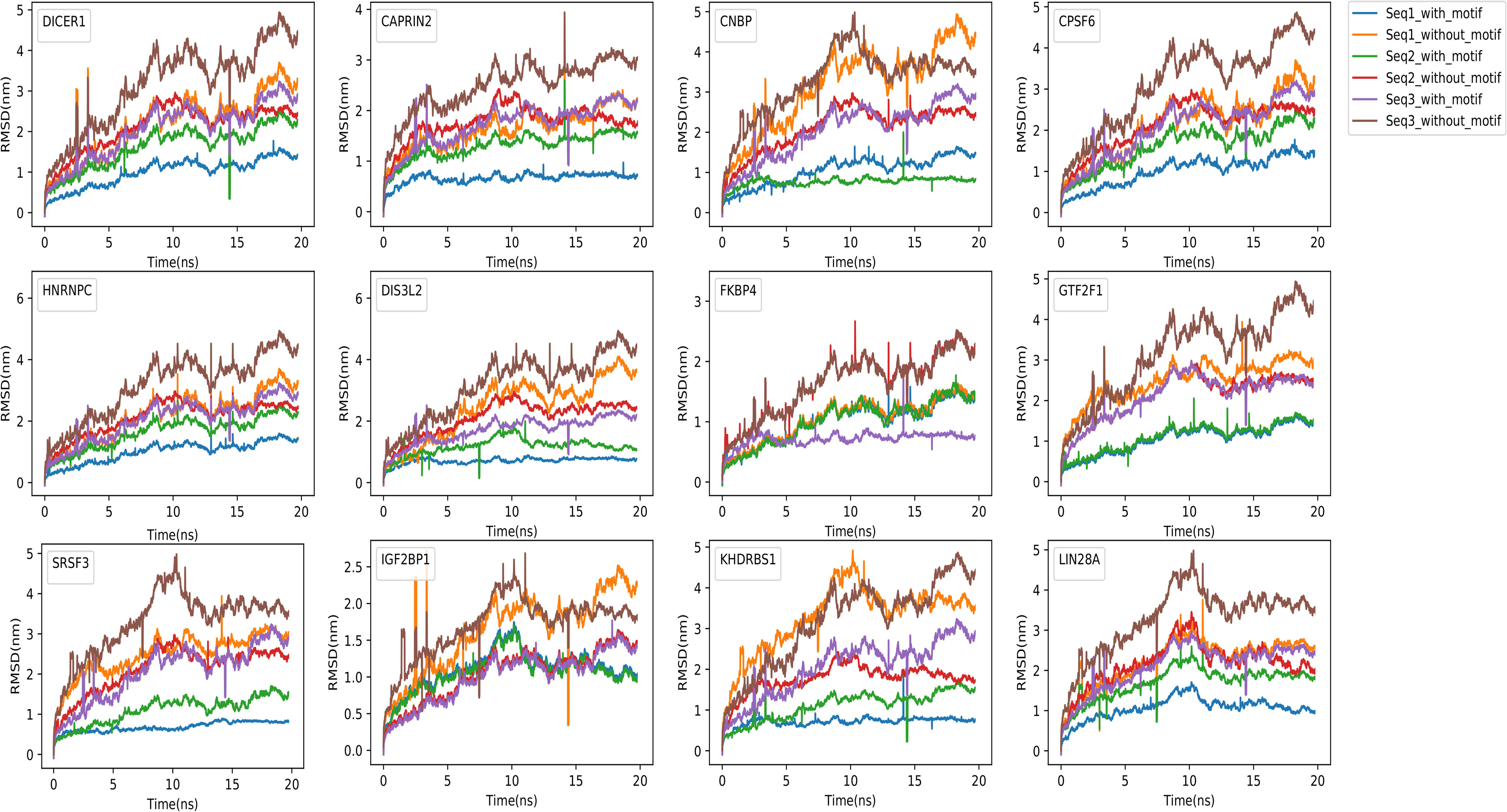
Comparative time dependent root mean square deviations (RMSD) plots for 12 different RBP-RNA complexes of with and without the prime motif. The trajectory was measured at 300 K for the 20-ns. Trajectory arcs for RBP-complex of three randomly selected RNA sequences with motifs are shown in blue, green and violet spike arcs whereas trajectory spike-arcs for RBP- complex without motif were shown in orange, red and brown color. The complexes with the prime motifs were found much stable than their counterparts without the prime motif.

For example, in case of AGO2, on comparative analysis of RMSD value of the AGO2-RNA first sequence complex with the prime motif, the value ranged from 0.1 nm to 0.7 nm and got stabilized at 0.5 nm whereas RMSD values for the complex without the motif ranged from 0.1 nm to 1.7 nm and got stabilized at 1.5 nm, which was less stable. Similarly the second pair with motif had RMSD ranging from 0.1 nm to 1.7 nm which got stabilized at 0.6 nm, whereas the same pair without the prime motif ranged had RMSD ranging from 0.3 nm to 1.4 nm and got stabilized at 1.4 nm. For the third pair, the AGO2-RNA complex of the third sequence with the prime motif showed deviation from 0.1 nm to 1.0 nm and got settled at 0.7 nm whereas the same sequence without the prime motif showed deviation from 0.0 nm to 2.0 nm and settled at 1.4 nm. In all the three cases of AGO-RNA complexes, the sequence with the prime motif was found to be more stable when compared to the one without the motif in the dynamic environment. Similar pattern was observed for all the 13 RBP and their triplicate pairs. Details can be found in Table 1.

**Table 1:** Table for RMSD value for selected 13 RBPs complexes with and without the prime motif. The identified prime motifs were found statistically enriched in the target sequences when compared to random regions. Molecular dynamics studies with and without these motifs clearly suggested their important role in binding where they were found responsible for stable complex formation between RBP and RNA.

| RBPs name | Sequence name | RMS Deviation range value with motif (nm) | Stablized_RMSD value with motif (nm) | RMS Deviation range value without motif (nm) | Stabilized RMSD value without motif (nm) |
| --- | --- | --- | --- | --- | --- |
| **AGO2** | RNA_seq1 | 0.1-0.7 | 0.5 | 0.1-1.7 | 1.5 |
|  | RNA_seq2 | 0.1-1.7 | 0.6 | 0.3-1.4 | 1.4 |
|  | RNA_seq3 | 0.1-1.0 | 0.7 | 0.1-2.0 | 1.4 |
| **CAPRIN2** | RNA_seq1 | 0.1-0.3 | 0.2 | 0.6-2.1 | 1.9 |
|  | RNA_seq2 | 0.3-1.8 | 1.3 | 0.7-2.2 | 1.5 |
|  | RNA_seq3 | 0.2-2.1 | 1.8 | 0.1-2.9 | 2.7 |
| **CNBP** | RNA_seq1 | 0.1-1.3 | 1.1 | 0.3-4.8 | 4.3 |
|  | RNA_seq2 | 0.3-1.8 | 0.5 | 0.3-2.5 | 2.0 |
|  | RNA_seq3 | 0.1-2.7 | 2.5 | 0.1-4.7 | 3.4 |
| **CPSF6** | RNA_seq1 | 0.3-1.2 | 1.0 | 0.3-3.2 | 3.0 |
|  | RNA_seq2 | 0.3-2.0 | 1.8 | 0.3-2.8 | 2.4 |
|  | RNA_seq3 | 0.1-2.9 | 2.7 | 0.1-4.8 | 4.1 |
| **DICER1** | RNA_seq1 | 0.3-1.3 | 1.0 | 0.3-3.5 | 2.9 |
|  | RNA_seq2 | 0.3-1.5 | 1.5 | 0.3-2.7 | 2.0 |
|  | RNA_seq3 | 0.1-2.5 | 2.3 | 0.1-4.8 | 4.2 |
| **DIS3L2** | RNA_seq1 | 0.1-0.3 | 0.3 | 0.3-2.5 | 2.5 |
|  | RNA_seq2 | 0.3-1.8 | 1.2 | 0.3-2.4 | 2.0 |
|  | RNA_seq3 | 0.1-2.3 | 1.8 | 0.1-4.2 | 4.0 |
| **FKBP4** | RNA_seq1 | 0.3-1.5 | 1.3 | 0.3-1.2 | 1.3 |
|  | RNA_seq2 | 0.1-1.7 | 1.1 | 0.1-2.5 | 2.2 |
|  | RNA_seq3 | 0.1-1.9 | 0.5 | 0.1-2.3 | 2.2 |
| **GTF2F1** | RNA_seq1 | 0.3-1.3 | 1.3 | 0.3-3.7 | 2.5 |
|  | RNA_seq2 | 0.3-1.9 | 1.7 | 0.5-2.5 | 2.0 |
|  | RNA_seq3 | 0.1-2.6 | 2.1 | 0.1-4.7 | 4.0 |
| **HNRNPC** | RNA_seq1 | 0.1-0.5 | 0.3 | 0.1-2.7 | 2.3 |
|  | RNA_seq2 | 0.1-2.3 | 1.8 | 0.1-2.3 | 2.0 |
|  | RNA_seq3 | 0.1-2.5 | 2.2 | 0.3-4.3 | 4.1 |
| **IGF2BP1** | RNA_seq1 | 0.3-1.7 | 0.9 | 0.3-2.6 | 2.1 |
|  | RNA_seq2 | 0.3-1.5 | 0.6 | 0.07-1.7 | 1.3 |
|  | RNA_seq3 | 0.1-1.7 | 1.5 | 0.1-2.6 | 1.6 |
| **KHDRBS1** | RNA_seq1 | 0.3-1.9 | 0.5 | 1.0-4.8 | 3.5 |
|  | RNA_seq2 | 0.3-1.2 | 1.2 | 0.3-2.1 | 1.2 |
|  | RNA_seq3 | 0.3-2.5 | 2.5 | 0.1-4.7 | 4.2 |
| **LIN28A** | RNA_seq1 | 0.3-1.4 | 0.7 | 0.3-3.0 | 2.2 |
|  | RNA_seq2 | 0.5-2.1 | 1.5 | 0.5-3.0 | 1.6 |
|  | RNA_seq3 | 0.1-2.2 | 2.2 | 0.1-4.9 | 3.1 |
| **SRSF3** | RNA_seq1 | 0.2-0.5 | 0.8 | 0.5-3.7 | 3.0 |
|  | RNA_seq2 | 0.3-1.2 | 1.2 | 0.5-2.5 | 2.1 |
|  | RNA_seq3 | 0.1-3.0 | 2.5 | 1.0-4.8 | 3.2 |

In the nutshell, the structural molecular dynamics study supported the identified binding spots for the RBP where it was clearly evident that the identified binding motif provided structural stability to the considered RBP-RNA complexes.

### Application: SARS-Cov2 genome was found to host RBP binding sites

Most of the deadly viruses are RNA viruses which exploit the host proteins to replicate, spread and survive. The best living example is nSARS-CoV-2. The emergence of the novel human corona-virus SARS-CoV-2 in Wuhan, China has caused a pandemic of respiratory disease (Covid19). The big scientific concern is that to this date very scarce and uncertain molecular information is available about the Covid19 patient’s molecular system as not much high-throughput studies have been carried out so far. There is almost absolutely no information on the host RBPs response during Covid19 infection despite of the fact that all such virus essentially require host RBPs to survive and replicate And RBP-RNA interaction studies hold prime importance in this regard also.

Therefore, we scanned the SARS-CoV-2 genome through RBPSpot to find the binding sites for RBPs which could have therapeutic value. Interestingly, out of 131 different model we found 22 different binding sites for 7 different RBPs (AIFM1 (2), BUD13 (3), CELF2 (4), RBM6 (3), UPF1 (2), TARBP2 (4) and KHSRP (4)) (Figure 11). Among these, AIFM1 interaction with viral polymerases in influenza virus infected cells is well studied (45). These all binding sites were found on anti-sense strand of the genome whose importance is for viral replication. During the infection, majority of immunoprecipitated RNA of Coronavirus were found originating from the anti-sense strand (46). Therefore, there is a possibility that these RBPs are helping in it’s transcription by binding to it’s negative strand. To check the stability of these RBPs with their binding site we also performed MD simulations study on two different sequence forms for each identified binding site (One with the binding site and another without it). Prior to this, we obtained complete 3D structures for AIFM1 and UPF1 from PDB and modeled the remaining five RBPs through homology modeling due to lack of complete defined structures for them. After modeling we evaluated the built 3D structure models using SAVES v6.0 (structure Activity validation server). Five RBPs PDB structures namely AIFM1, BUD13, CELF2, TARBP2 and UPF1 passed through verification filter like PROCHECK and WHATCHECK except KHSRP and RBM6. When we analyzed the model structure for KHSRP and RBM6 with both program it gives 80.9% and 83.5% residues in allowed regions in the Ramachandran plot but for good quality model, over 90% residues are expected in the most favored region and lack of loop filtering causing side-chain packing inaccuracies. Subsequently, on analyzing the RMSD graph (Supplementary Figure 2) for all the seven RBPs it was found that that five out of seven RBP-RNA complexes were stable with prime motif compared to the RBP-RNA complexes counterpart without the main motif. This part of the study was done just to showcase the application of the developed approach. The finding made in this section may be used for further study for Covid research groups.

**Figure 11:**
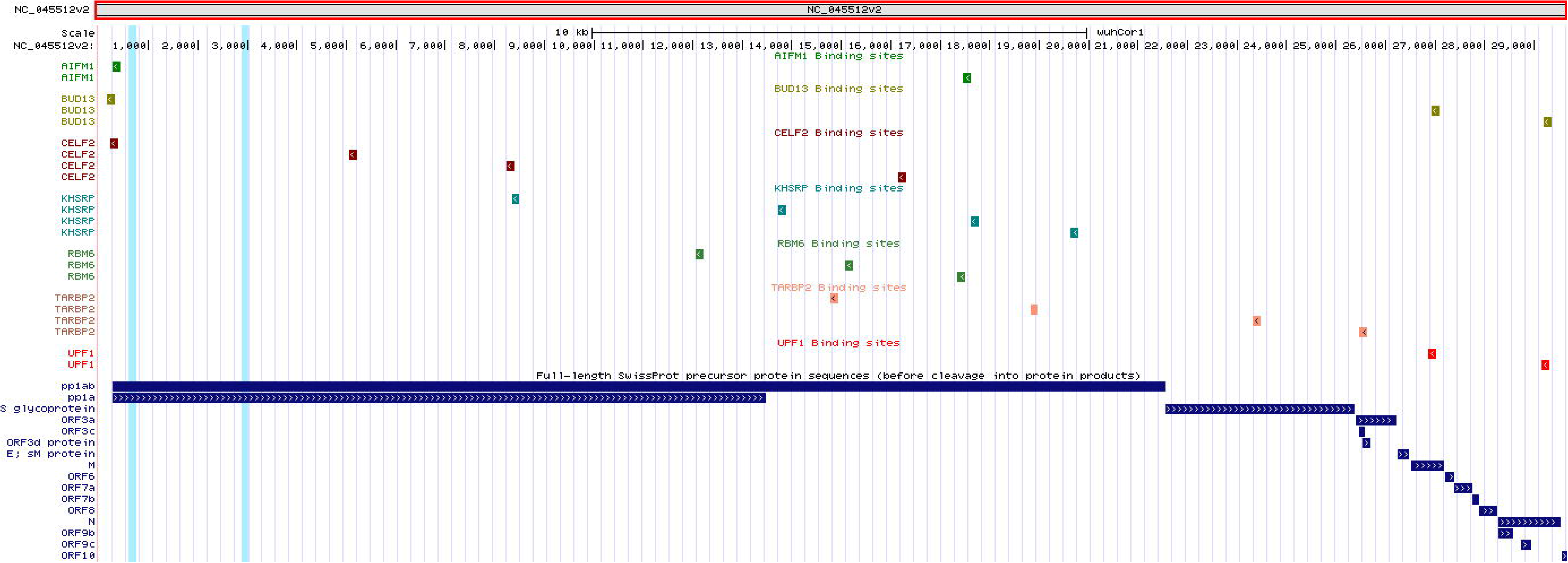
Application of RBPSpot reports the binding sites for seven different RBP on nSARS- CoV2 genome. A total for 22 such binding sites were discovered across the nSARS-CoV2 genome, all which existed across the negative strand of the virus genome.

## Conclusion

A living system is a continuous outcome of the regulatory setup working for that system in the background. RNA binding proteins define one such critical regulatory component of the system which is present at almost every post-transcriptional regulatory event but about which our understanding is still nascent and evolving. How they select their targets and carry out interactions in functional manner is largely ambiguous. With the advent of high-throughput techniques like CLIP-seq and interactome capture, the information on genes recognized as RBPs and their interactions are growing continuously. Such high-throughput data on interactions are very valuable resources to construct the interaction models. The present study used the same from CLIP-seq experiments. However, there are several other critical factors involved which are required to be build these interactions models with high accuracy. This involves proper negative data-sets screening, appropriate motif discovery strategy, and contextual information derivation. All of them are interconnected with each other and success of any such RBP binding site discovery tool depends highly on this. Without proper datasets, correct binding specific motif candidates are hard to be found. The motif finding step itself needs to consider the sparse nature of RBP binding sites and need to anchor correctly so that correct surrounding could be recognized to provide the contextual information. Otherwise, such motifs occur frequently even in the non-binding regions, and wrong context may easily compromise the accuracy. When all these information are applied through effective machine learning algorithms, consistently high level accuracy is achievable. It was comprehensively and comparatively benchmarked against some recent tools where it outperformed them consistently across a wide number of datasets and RBPs. It also showcased that when a DNN is trained properly on suitable properties with appropriate biological insights, the developed system could easily outperform much complex deep-learning based approaches where such learning is done through automated feature extraction process using complex layers like CNN and LSTM etc. Such complex deep learning approach may be suitable for unstructure data where features could not be identified easily. However, when features are identifiable and structured, simpler machine learning approaches can outperform them easily. The developed approach in this study, RBPSpot, can identify the binding sites of existing RBPs in human system as well as it becomes one of few tools where users can put their own data and raise their own models for any species and any RBP. The software is freely available as a webserver as well as as an standalone program.

From here, we visualize that incorporation of spatio-temporal and other interactome network information for RBPs as the another dimension to explore to further improve our understanding on RBP-RNA interactions. This is something which still remains largely unaddressed. Some encouraging recent developments have happened (47, 48) which promise that incorporation of back- end network and interaction information on RBP RNA interactions could add more value towards recognition of functional and dynamic nature of RBP RNA interactions which could further boost interaction spot identification process. Also, the findings made here from the contextual information like CG enrichment in the flanking regions must be explored further for their functional roles associated with such binding sites. RNA modifications on CG and likewise other important contextually important factors found in this study may further provide reasoning for spatio-temporal nature of these interactions which would mark another level of development in our understanding towards RBP RNA interactions and regulation.

## Declarations

### Availability of data and materials

All the secondary data used in this study were publicly available and their due references and sources have been provided. All data and information generated/used, methodology related details etc have also been been made available in the supplementary data files provided along with and also made available through the related open access server at https://scbb.ihbt.res.in/RBPSpot/. The software has also been made available at Github at:https://github.com/SCBB-LAB/RBPSpot

### Authors’ contributions

NKS and SG carried out the computational part and benchmarking of the study. PK developed the FM-Index and BWT based inexact k-mer search script. AK carried out the structural analysis and molecular dynamics simulation. UKP helped in statistical analysis. RS conceptualized, designed, analyzed and supervised the entire study. NKS, SG, AK and RS wrote the MS.

## Acknowledgments

We are thankful to the Director, CSIR-IHBT, for his kind support. We are thankful to Dr. Indu Gangwar for her inputs for the study. We are thankful to DBT for the funding support they gave for this project. NKS is thankful to CSIR for financial support as project associateship. UKP and PK are thankful to ICAR, New Delhi for providing support in Ph.D. UKP, NKS, and PK are thankful to Academy of Scientific and Innovative Research (AcSIR).

## Ethics approval and consent to participate

Not applicable.

## Consent for publication

Not applicable.

