## supplementary methods for "RBPSpot: Learning on Appropriate Contextual Information for RBP Binding Sites Discovery"

**Supplementary method:**

Burrow-Wheeler Transformation and FM-index based search algorithm is given below:

A sequence *“S”* of length len(s) is made of from a set of alphabet letters *Φ* = {A, C, G, U}. S*[i]* denotes the *i*-th position of base as (0<=i<=len(s)-1) and S*[i,j]* represents a subsequence which starts at position *i* and ends at *j*. This way each such sequence region of length len(S) generates len(S)-6 k-mer subsequences of length 6, which is converted into a set of unique 6-mers. Initially all of them are assumed to be a potential candidate which could given birth to a motif seed, which becomes clear after continuous search across all CLIP-seq peak regions for every such candidate and enrichment analysis for over-representation. Mapping each of such subsequence query on reference sequence *“S”* (CLIP-seq peak regions) required to do two basic operations: 1) find the frequency of the query subsequence “*q”* in the reference sequences, and 2) locate the position the subsequence *“q”* in the reference sequence “S”.

Step 1: Input: k-mers set as set of queries “q” and Set of reference sequences “S_N_”.

Step 2. Perform BWT transformation of S_N._

Step 3. Make the dictionary of count of all bases by using BWT and used to find Tally Dictionary.

Step 4. Make tally matrix using FM index base search of k-mer.

Step 5. Using BWT transformed S_N_ set of reference sequences, generate the base count dictionary and tally dictionary. Tally dictionary stores the base position records.

Step 6: Repeat:

Step 7: Keep updating the Tally dictionary according to the found base positions.

Step 8: Until the cycle reaches the end reference sequence, for all elements of S_N_.

Step 9: Generate the final first column dictionary holding base and position in sorted manner.

Step 10: Search query *k-mer* *“q”* in the reference set using first column dictionary and tally dictionary.

Step 11: Return: Report all found locations.

The inexact search is done further by generating all possible k-mers with given mismatch level to the root k-mer. These similar relatives are parallely searched across the generated indices through above mentioned steps.

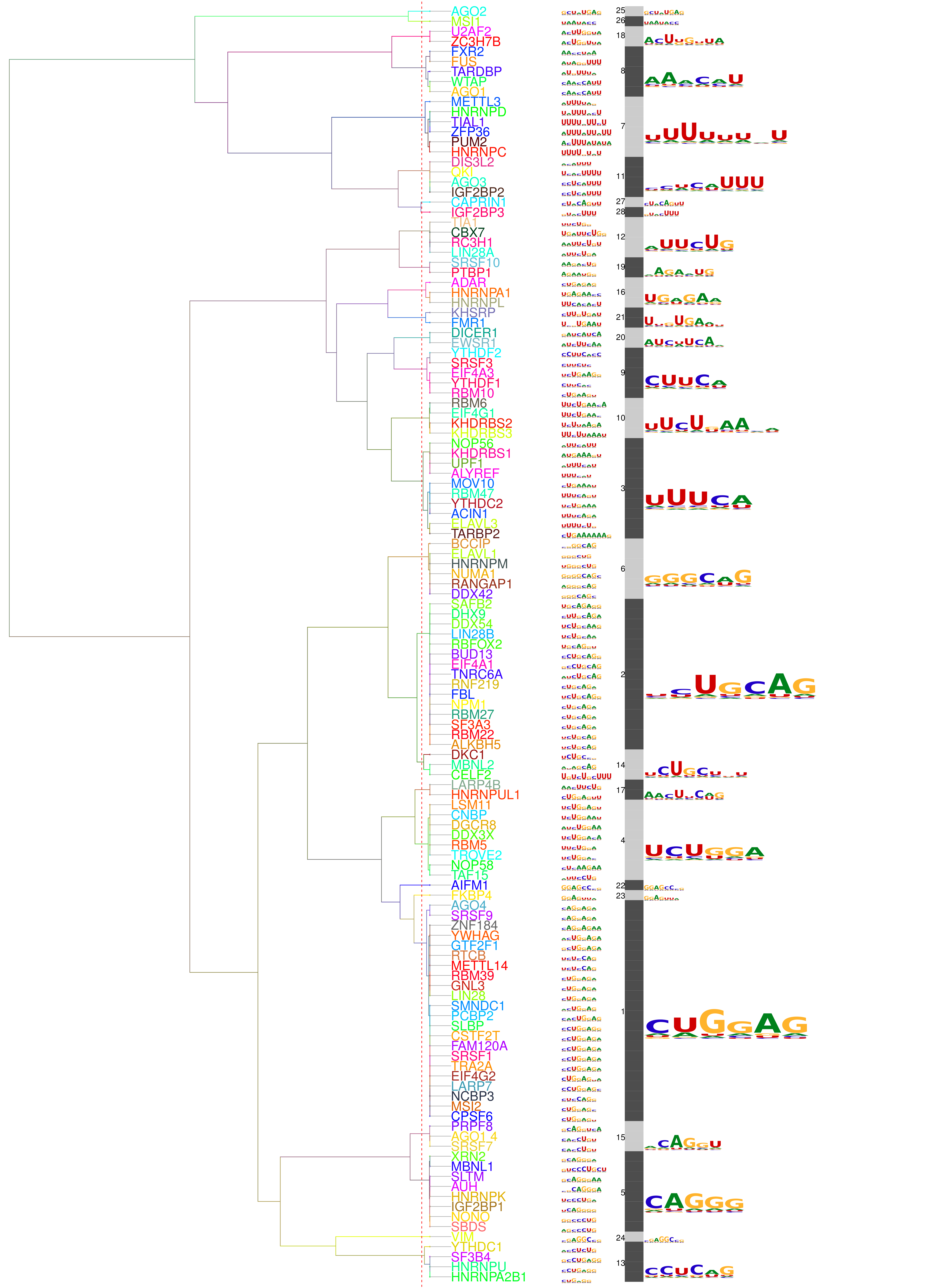
**Supplementary Figure:**

**Supplementary Figure 1**: Motif clustering of the RBPs. A total of 28 different clusters for 127 different RBP were formed. The members of these clusters shared high similarity among their binding site prime motifs. However, despite of sharing close similarity, their binding were found highly contextual.

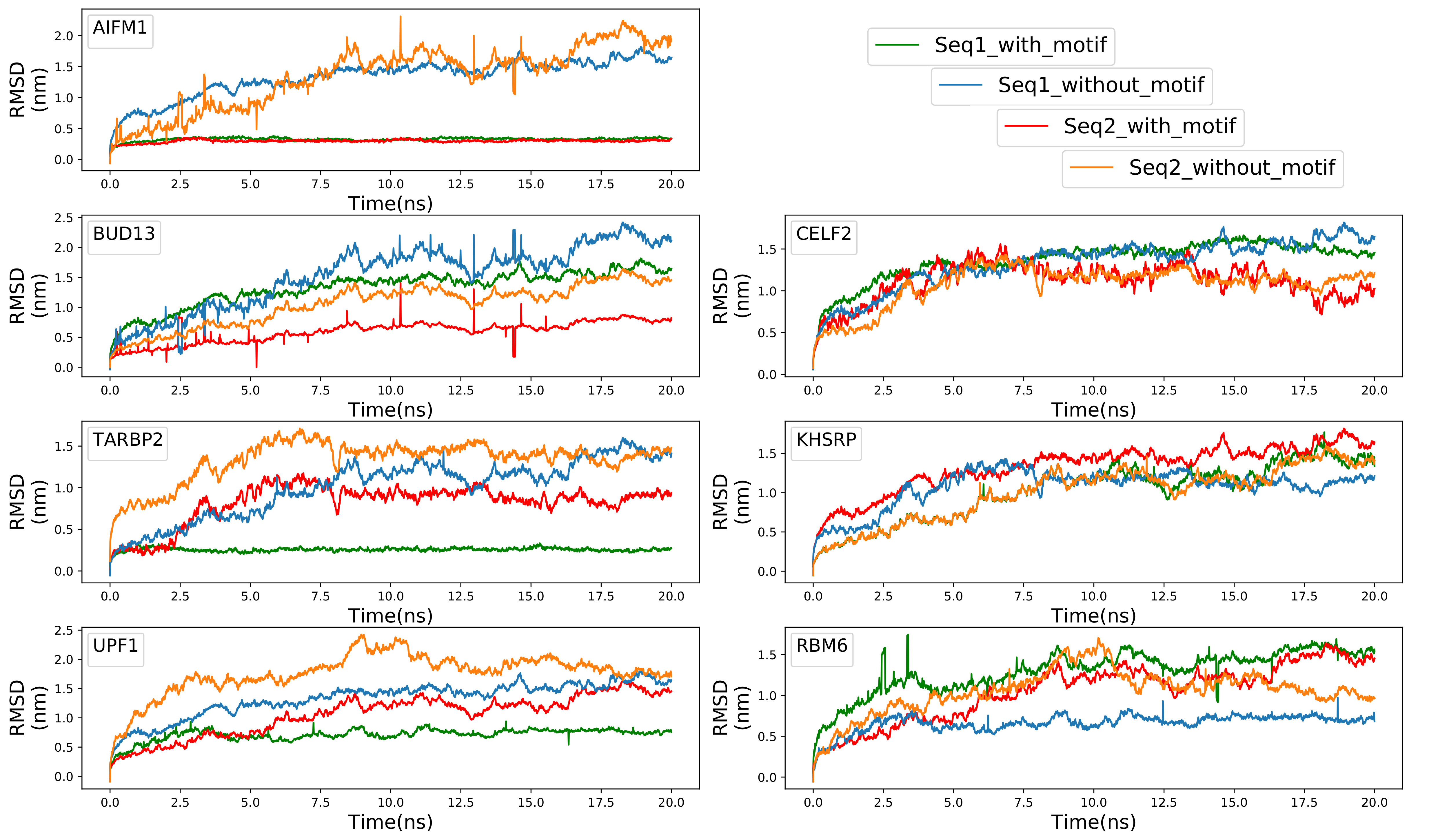

**Supplementary Figure 2**: **Comparative time dependent root mean square deviations (RMSD) plots for the seven diffrent RBP-RNA complexes for interaction with nSARS-CoV-2 genome. It was measured with and without the main motif. The trajectory was measured at 300 K for the 20-ns. The molecular dynamics study supported most of the discovered binding sites identified by RBPSpot.**
